# A massively parallel synthetic gene atlas for learning compact *cis*-regulatory grammar across cellular contexts

**DOI:** 10.64898/2026.09.13.751267

**Authors:** Timo Hagen, Isabelle Liu, Ariana Nagainis, Noosha Khosrojerdi, Hassan Yousefi, Tess Gunnels, Amir Moarefi, Kevin Green, Nazli Azimi, Amir Momen-Roknabadi, Kevin R Kipp, Hani Goodarzi

**Author notes:** These authors contributed equally (Timo Hagen, Isabelle Liu, Ariana Nagainis). Correspondence: Amir Momen-Roknabadi, Kevin R Kipp, Hani Goodarzi.

## Abstract

Virtual-cell models increasingly learn from large perturbation atlases, but their view of *cis* regulation remains limited to endogenous genes embedded in broad native regulatory contexts. Here we introduce Therna Biosciences’ Chronos platform and its first public dataset release, comprising two complementary modules of a massively parallel synthetic gene atlas: *Penta-47×27K* for 5′ UTRs/internal-promoter elements and *Tria-47×28K* for 3′ UTR stability elements. Together, these datasets measure ∼60,000 compact *cis*-regulatory elements across approximately 50 cell lines in one pooled experiment. The choice of cell lines here, provided to us courtesy of Tahoe Tx, was rooted in our aim to make this dataset maximally useful for the virtual-cell modeling community. At Therna, we routinely apply Chronos — part of our RNA-Logix™ platform — beyond cancer cell-line pools, to systems such as primary cells and organoids, via LNP-formulated mRNA delivery.

Chronos captures both transcriptional and post-transcriptional gene expression control. Episomal DNA delivery measures DNA-normalized mRNA output, while direct RNA delivery with longitudinal sampling reliably quantifies RNA decay. The RNA-delivery arm uses chemically modified synthetic mRNA incorporating N1-methylpseudouridine, a modification widely used in mRNA therapeutics. To resolve high-complexity libraries of up to 30,000 elements, Chronos pushes the sensitivity limits of single-cell RNA sequencing to quantify individual RNA molecules within single cells. By massively expanding gene regulatory networks with synthetic genes whose variable regulatory code is short and defined, Chronos provides an auxiliary *cis*-regulatory lens for virtual-cell modeling and enables context-specific *cis*-*trans* regulatory interactions to be learned directly.

## Introduction

Building predictive models of the cell requires data that systematically connect perturbations to cellular responses. Large-scale single-cell perturbation studies measure how cells respond to genetic perturbations, chemical perturbations, and combinations of perturbations across many cellular contexts. *Tahoe-100M*^1^, Perturb-seq-style CRISPR perturbation resources^2–5^, and related perturbational datasets have helped define a practical experimental format for training and evaluating virtual-cell models^6–9^: perturb the cell, measure the transcriptome, and learn how cellular state changes as a function of context.

Because these datasets systematically vary *trans*-acting perturbations while leaving the underlying *cis*-regulatory sequences largely fixed, the models trained on them inherit the same emphasis on *trans*-regulatory responses. Transcriptome-response models such as Arc’s STATE^10^ and STACK^11^, Xaira’s X-Cell^12^, Tahoe’s Rhaister^13^, and related virtual-cell approaches^6,7,9^ predict how gene expression changes after perturbation or across cellular contexts. Chromatin and 3D-genome models, such as Cleopatra^14^, and single-cell epigenomic models, such as EpiFoundation^15^, address *cis*-regulatory biology more directly; however, the datasets needed to train these models remain sparse and costly to generate at scale.

Consequently, existing models are well suited to predicting how cells respond when signaling pathways, drug targets, transcription factors, chromatin regulators, or other cellular effectors are perturbed, but they do not densely sample *cis*-regulatory sequence space. With rare exceptions for naturally occurring variants, each cell contains the same endogenous genome, and models observe regulatory sequences through the limited set of endogenous loci represented in the training data. This creates a fundamental asymmetry: the number of measured genes is on the order of 20,000, but the sequence contexts around endogenous genes is vast. While a faithful model of gene-expression control must learn both *cis*-regulatory sequence grammar and the *trans*-regulatory contexts in which that grammar is interpreted, single cell transcriptomic data does not provide sufficient data scale to accomplish this task, i.e. link the expression of genes to specific sequence elements in surrounding enhancers, promoters, and untranslated regions.

Gene expression is governed by transcriptional and post-transcriptional regulatory programs^16–18^. Transcriptional programs determine how much RNA is produced from a DNA template, whereas post-transcriptional programs determine how RNA is processed, translated, stabilized, or degraded after it is made. These layers are coupled in endogenous genes and are difficult to separate because each locus carries a large native regulatory architecture. We reasoned that this problem becomes more tractable if the measured gene space is expanded with synthetic genes. Therna Biosciences’ Chronos platform is designed to supply the missing axis by compacting *cis*-regulatory grammar into synthetic gene libraries that can be read out through both DNA- and RNA-delivery formats.

In our DNA-delivered format, we varied the 5′ UTR downstream of a constant core promoter to study internal *cis*-regulatory elements. Other massively parallel assays use this position differently — not as the tested element, but to tag a candidate upstream of, or overlapping, the transcription start site. lentiMPRA^19^ barcodes a candidate located upstream of a constant minimal promoter. STAP-seq^20^ instead places candidates at the minimal-promoter position, downstream of a constant enhancer, and reads the transcript’s 5′ end to map them.

Yet the 5′ UTR is not just a convenient tagging site. Sequences embedded within a transcribed 5′ UTR can themselves carry transcriptional and post-transcriptional regulatory information. Patient-derived prostate cancer mutations, for example, have been shown to create promoter-like elements that alter transcript levels^21^. To study this at scale, we designed a synthetic library of ∼30,000 elements within the 5′ UTR, profiled across 47 cellular contexts, the most comprehensive set of 5′ UTR/internal-promoter regulatory elements to date.

Therna Biosciences is publicly releasing two foundational Chronos datasets generated in a 48-cell-line Mosaic cancer cell-line pool (courtesy of the Tahoe Tx team): *Penta-47×27K*, measuring transcriptional activity of 5′ UTR/internal-promoter regulatory elements, and *Tria-47×28K*, measuring the impact of 3′ UTR elements on mRNA stability. Together, these datasets profile compact regulatory elements across 47 cellular contexts and align the resulting *cis*-regulatory atlas to *Tahoe-100M*. This design makes Chronos a complementary *cis*-regulatory layer for perturbation atlases and virtual-cell models, which otherwise remain largely anchored to endogenous transcriptome responses.

## Results

### Chronos compacts *cis*-regulatory grammar while preserving cell-context resolution

We designed Chronos to expand the number of measured genes while constraining the regulatory code that varies between them. Each synthetic gene contains a shared architecture and a compact variable regulatory element. In the DNA-encoded 5′ UTR library, the variable element occupies a compact window within the 5′ UTR, downstream of a constant EF1α core promoter. Most existing datasets use this region only to encode a barcode tied to a variable promoter region. Here, we instead made it the variable *cis*-regulatory element itself, providing the first comprehensive characterization of internal elements and their role in gene expression. In the RNA-encoded 3′ UTR library, the variable element occupies a compact window within the 3′ UTR, downstream of a constant coding sequence and upstream of the poly(A) tail (**Fig. 1a**). This design creates tens of thousands of additional measured genes whose regulatory grammar is largely defined by variations in a short sequence window rather than by the full endogenous locus.

**Fig. 1.**
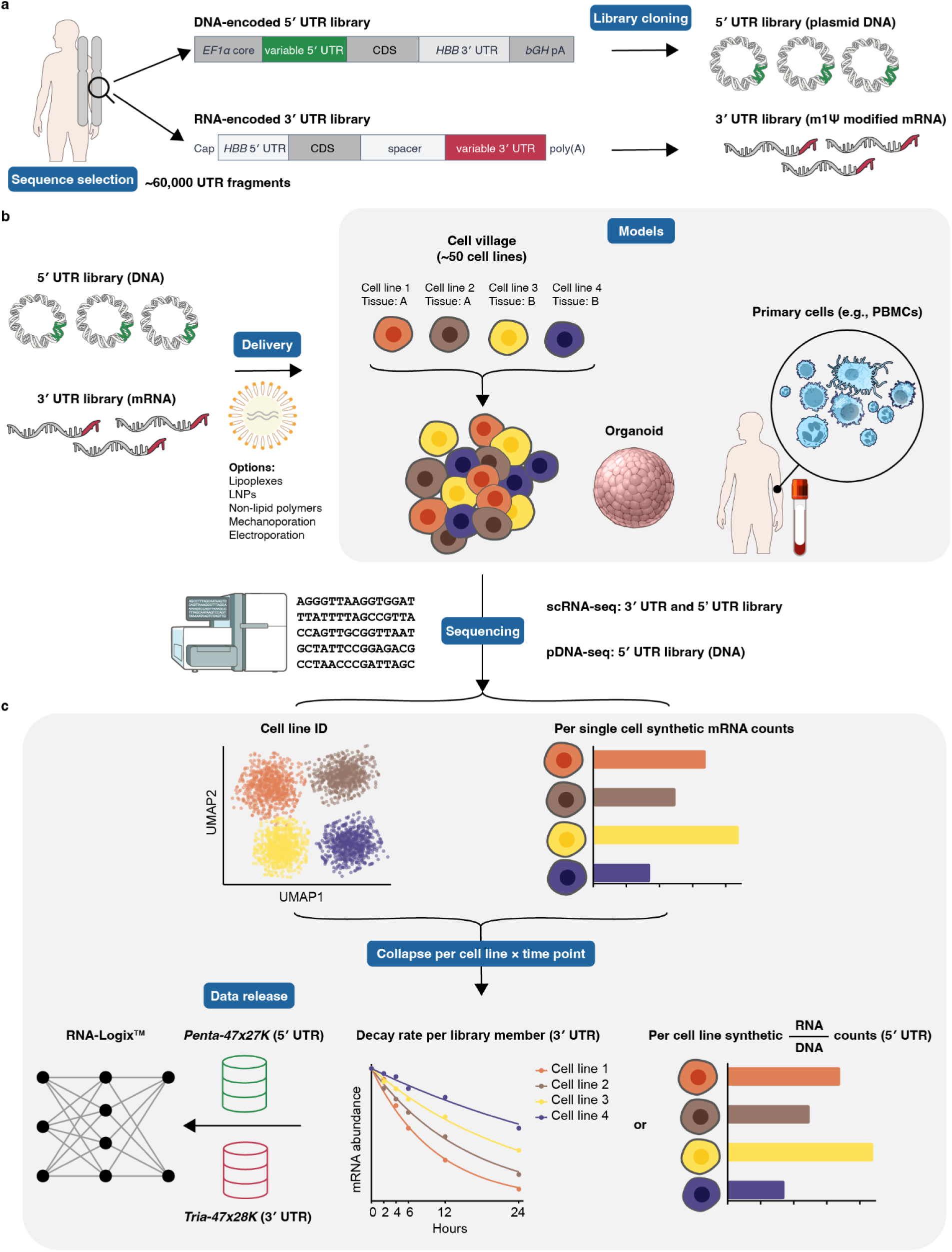
Chronos platform: the experimental and informatics workflow used to generate *Penta-47×27K* and *Tria-47×28K*. **a**, Library design. Compact synthetic gene libraries were generated by sampling ∼30,000 UTR fragments each from 5′ UTR and 3′ UTR regions of the human transcriptome. The 5′ UTR library (green) is delivered in episomal/plasmid DNA format, with transcription driven by an EF1α core promoter, and comprises 50-nt variable fragments. The 3′ UTR library (red) is delivered as synthetic mRNA generated by T7 polymerase *in vitro* transcription with Clean-Cap-AG and N1-methylpseudouridine (m1Ψ), and comprises 240-nt variable fragments. **b**, Experimental schematic. Libraries are delivered by the method matched to each regulatory question: episomal/plasmid DNA delivery yields DNA-normalized expression output, while direct RNA delivery with longitudinal sampling yields decay-rate estimates. Both formats can be applied across a range of cellular models, including pooled cell-line villages, organoids, and primary cells (e.g., PBMCs). This release uses the Tahoe Mosaic cancer cell-line pool from which *Tahoe-100M*^1^ was generated, enabling pooled measurements across 47 cell-line contexts in a single experiment (four cell lines shown for simplicity). **c**, Data analysis. Whole-transcriptome profiles are used for single-cell context assignment, after which synthetic-gene reads are collapsed to the context level to produce final expression and decay-rate matrices ready for downstream model training.

The two delivery formats isolate distinct regulatory layers, allowing Chronos to capture both transcriptional and post-transcriptional regulatory programs. In the episomal/plasmid format, compact *cis*-regulatory fragments are delivered as DNA templates, and the resulting RNA abundance is measured in each cellular context (**Fig. 1b**). RNA counts are normalized to the DNA counts of each fragment in the input pool, producing a DNA-normalized expression estimate that reflects RNA output per input DNA molecule (**Fig. 1c**). In the RNA-delivery format, synthetic RNAs (modified or unmodified) are delivered directly and sampled across time (**Fig. 1b**). This format removes transcription from the measurement and enables quantification of post-transcriptional regulation through RNA persistence over time, and decay rate estimates (**Fig. 1c**). Together, the DNA and RNA formats allow compact *cis*-regulatory elements to be decomposed into transcriptional and post-transcriptional outputs in the same cellular frame.

Running both libraries in a pooled cell system measures the same compact regulatory variants across approximately 50 cellular contexts in one experiment (**Fig. 1c**). Because each cell contains only a subset of the synthetic library, the output is not interpreted at the individual-cell level. Instead, Chronos collapses synthetic-gene measurements across cells within each assigned context, producing cell-line- or cell-type-level matrices of regulatory output (**Fig. 1c**). This design preserves the scaling advantage of single-cell profiling while producing the context-resolved measurements needed for sequence-to-function modeling.

Specific implementations in *Penta-47×27K and Tria-47×28K* include compact 5′ UTR designs with an internal promoter architecture and compact 3′ UTR RNA-stability designs. The 5′ UTR design comprises 30,000 distinct 50-nucleotide sequences sampled from the human transcriptome. The 3′ UTR design comprises 30,000 synthetic 240-nucleotide sequences sampled from the human transcriptome. *Penta-47×27K* and *Tria-47×28K* represent the most extensive cell-context datasets to date for DNA- and mRNA-delivered regulatory elements, respectively, each profiled across 47 matched cellular contexts. These libraries should be interpreted as implementations of the broader Chronos principle: compress the variable *cis*-regulatory code into a learnable window, measure regulatory output across many cellular contexts, and use the paired cell-context information to learn *cis*-*trans* interactions.

### Chronos maps compact regulatory output across the Tahoe Mosaic cell-line pool

For this study, the Tahoe team kindly provided access to the Mosaic cancer cell-line pool used to generate *Tahoe-100M*^1^, which served as the cellular substrate for Chronos. This enables Chronos to be joined to a widely used perturbation atlas. However, the Chronos design is not specific to cell-line villages and can be applied to other pooled or complex cellular models, including primary cells, organoid cultures, iPSC-derived cell types, and *in vivo* models as well as to LNP-formulated mRNA delivery.

The pooled Mosaic format is central to *Penta-47×27K and Tria-47×28K*. Rather than profiling each cell line separately, Chronos quantifies gene expression dynamics for the same synthetic regulatory library across the pooled panel in parallel. 48 cell lines were transfected, of which 47 were confidently distinguished and assigned by transcriptome-wide label transfer; the complete assay contains these 47 cellular contexts, with analysis-specific QC applied to remove outlier cells and genes. Whole-transcriptome profiles identify each cell’s context, and targeted synthetic-gene reads quantify the synthetic library. The resulting data are collapsed by cell-line context to produce matched regulatory-output matrices across the pool (**Fig. 1c**).

To profile RNA decay, the direct RNA delivery format was processed using Parse split-pipe processing, NVIDIA rapids-singlecell^22^ analysis, and scANVI-based^23^ label transfer from matched cell-line references in *Tahoe-100M*. This assignment strategy enabled Chronos measurements to be aligned to the same cell-line identities used in the broader *Tahoe-100M* resource (**Fig. 2a**). Similarly, to evaluate 5’ *cis*-regulatory control, the episomal plasmid format was processed using an internally developed workflow, including 10x Genomics cellranger^24^ processing, NVIDIA rapids-singlecell analysis, and scANVI-based label transfer from matched cell-line references in the RNA delivery library. We recovered reporter-positive cells across all 47 cellular contexts and all sampled time points in both *Penta-47×27K and Tria-47×28K*, although recovery varied substantially among cell lines (**Extended Data Fig. 1**), demonstrating broad representation of the pooled panel while motivating cell-line-aware filtering and normalization (**Fig. 2b**).

**Fig. 2.**
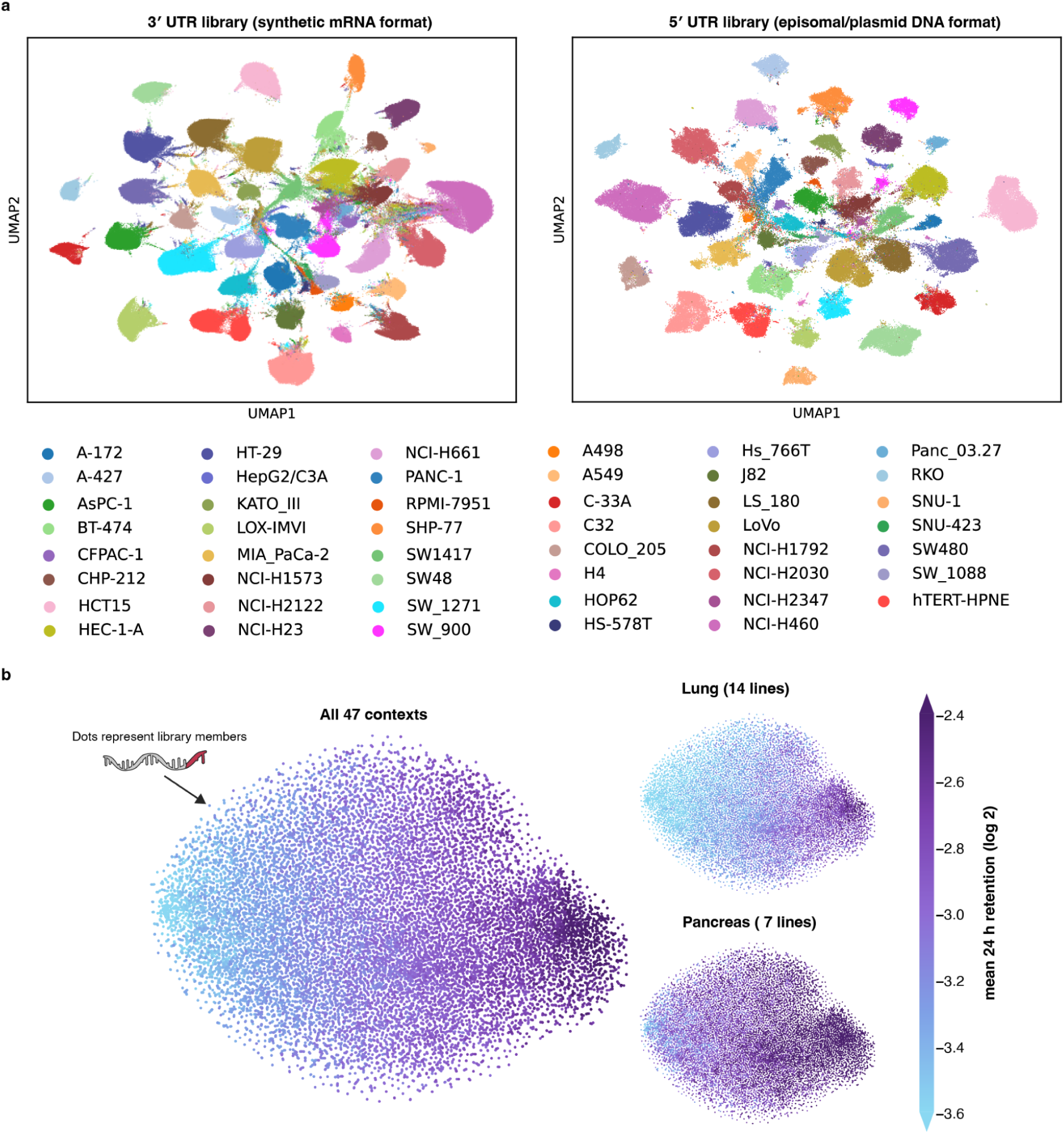
The pooled cell-line panel resolves both cell identity and tissue-specific activity. **a**, Cell-line identities recovered across the pooled reporter assays. UMAP representations of single-cell transcriptomes from the pooled 47-cell-line panel after assignment of cells to their corresponding cell-line identities. Shown are cells recovered from the 3′ UTR library delivered as synthetic mRNA (left) and the 5′ UTR library delivered as episomal plasmid DNA (right). Each point represents one cell, and colors denote assigned cell lines. The separation of the expected cell-line populations confirms broad recovery and reliable cell-identity assignment in both delivery formats. **b,** t-SNE of synthetic mRNA library members (each point, one 3′ UTR variant) embedded from their 24-h RNA-retention profiles across cellular contexts. Left, all 47 cell-line contexts; right, the lung (top, 14 lines) and pancreas (bottom, 7 lines) subsets. Points are colored by mean 24-h retention (log_2_), low (light) to high (dark). Members form a continuous stability gradient rather than discrete clusters, and the tissue subsets redistribute members while preserving this global axis, indicating that RNA stability is context-dependent.

### Chronos recovers spike-in standards, known stability elements, and known transcription-factor motifs

We next evaluated whether Chronos measurements behaved as expected using internal controls and orthogonal in-house datasets. The RNA-delivery libraries included concentration spike-ins spanning known relative abundance levels, as well as positive and negative stability controls. These controls allow sample-level recovery, normalization, and stability ranking to be evaluated independently of the learned sequence models.

The concentration spike-ins recovered the expected abundance structure after normalization, supporting the use of sample-level scaling factors for comparing synthetic mRNA counts across time points and conditions (**Fig. 3a**). Stability controls showed the expected separation between stable elements and destabilized mutants, confirming that Chronos can recover known post-transcriptional effects in pooled cellular contexts (**Fig. 3b**).

**Fig. 3.**
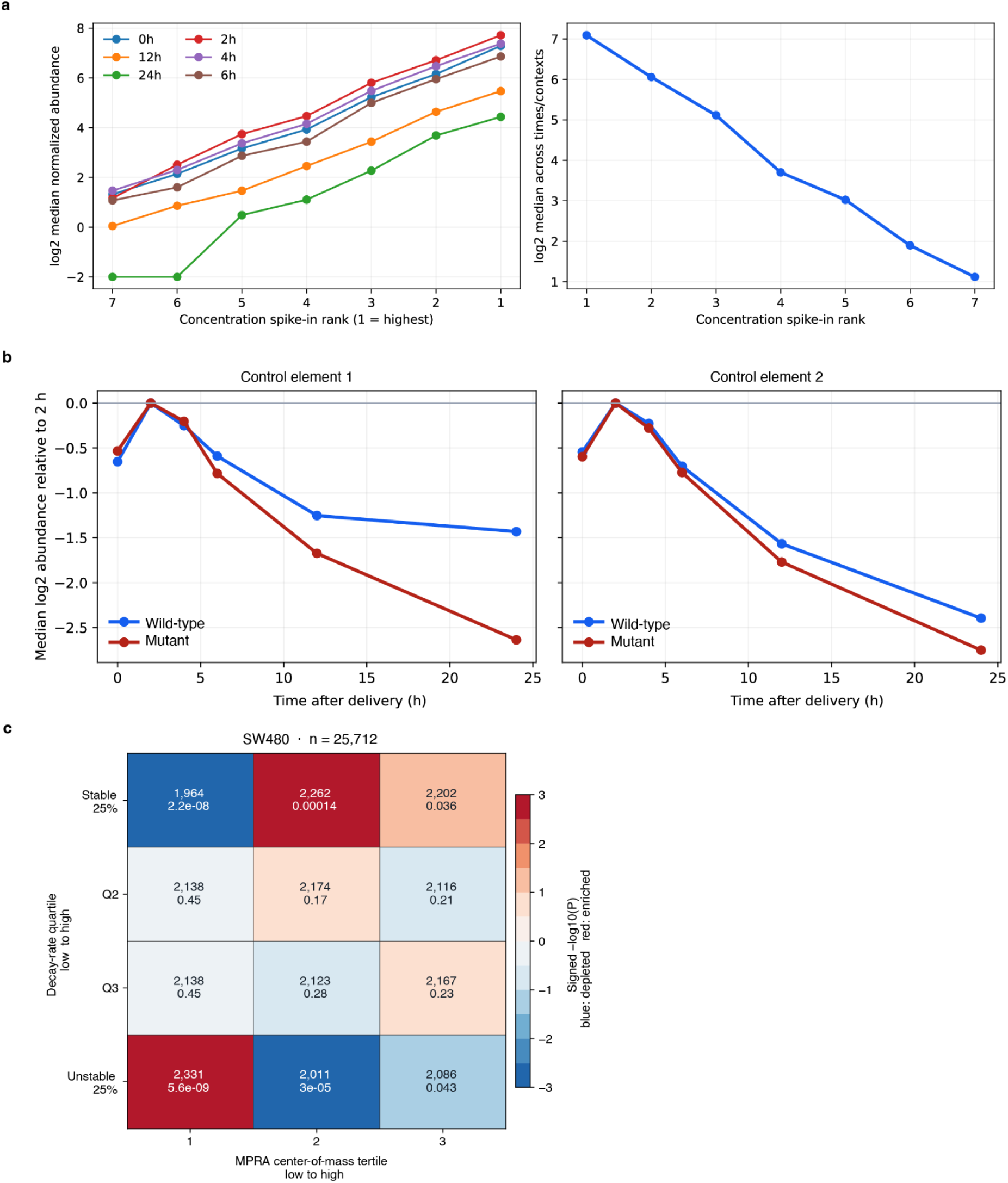
Technical and biological validation of the 3′ UTR stability assay. **a**, Recovery of a seven-member concentration spike-in series across the six sampled time points. Median normalized abundance increased in line with the expected spike-in concentration at each time point (left). Median abundance across cell lines and time points showed a perfectly monotonic relationship with the dilution rank (right; Spearman ρ = −1.00, where rank 1 denotes the highest concentration). **b**, Median abundance trajectories for two previously characterized RNA-stability elements and their destabilizing point mutants, normalized to abundance at 2 h. Control element 1 and its less stable variant are shown on the left; control element 2 and its less stable variant are shown on the right. In both cases, the destabilizing point mutant declined more rapidly than its corresponding control element, recovering the expected direction of the point-mutation effects. **c**, Comparison of 3′ UTR decay rates with an independent MPRA measurement in SW480 cells (n = 25,712 elements). Elements were divided into decay-rate quartiles, from stable to unstable, and MPRA center-of-mass tertiles, from low to high reporter output. Each cell reports the number of elements and the two-sided Fisher exact-test q value after Benjamini–Hochberg correction across the 12 cells in that panel. Color represents signed −log10(P): red indicates enrichment and blue indicates depletion. Rapidly decaying elements were enriched in the lowest MPRA-output tertile, whereas stable elements were depleted from this tertile, providing independent directional support for the measured stability differences.

We also compared Chronos 3′ UTR regulatory outputs against existing in-house MPRA datasets measuring post-transcriptional activity, an assay format also used in published platforms^25^. These analyses test whether compact *cis*-regulatory elements that behave similarly in conventional reporter assays also show concordant activity in Chronos, and whether cell-context-specific Chronos effects align with prior measurements of post-transcriptional regulatory activity. These evaluations provide an orthogonal check that Chronos is not only technically reproducible, but biologically calibrated to established *cis*-regulatory measurements (**Fig. 3c**).

Next, we evaluated whether the transcriptional (5′ UTR/episomal) arm showed similar internal reproducibility and biological grounding. Panel-level element detection was strongly correlated between the 24 h and 36 h time points (Spearman ρ = 0.816, Pearson r = 0.796). Across all 47 matched cell lines, within-line 24-36 h Spearman correlations had a median of 0.242 and ranged from 0.011 to 0.583. Thus, the experiment showed strong panel-level reproducibility, while the persistence of element-level signal between time points varied across cellular contexts (**Fig. 4a**).

**Fig. 4.**
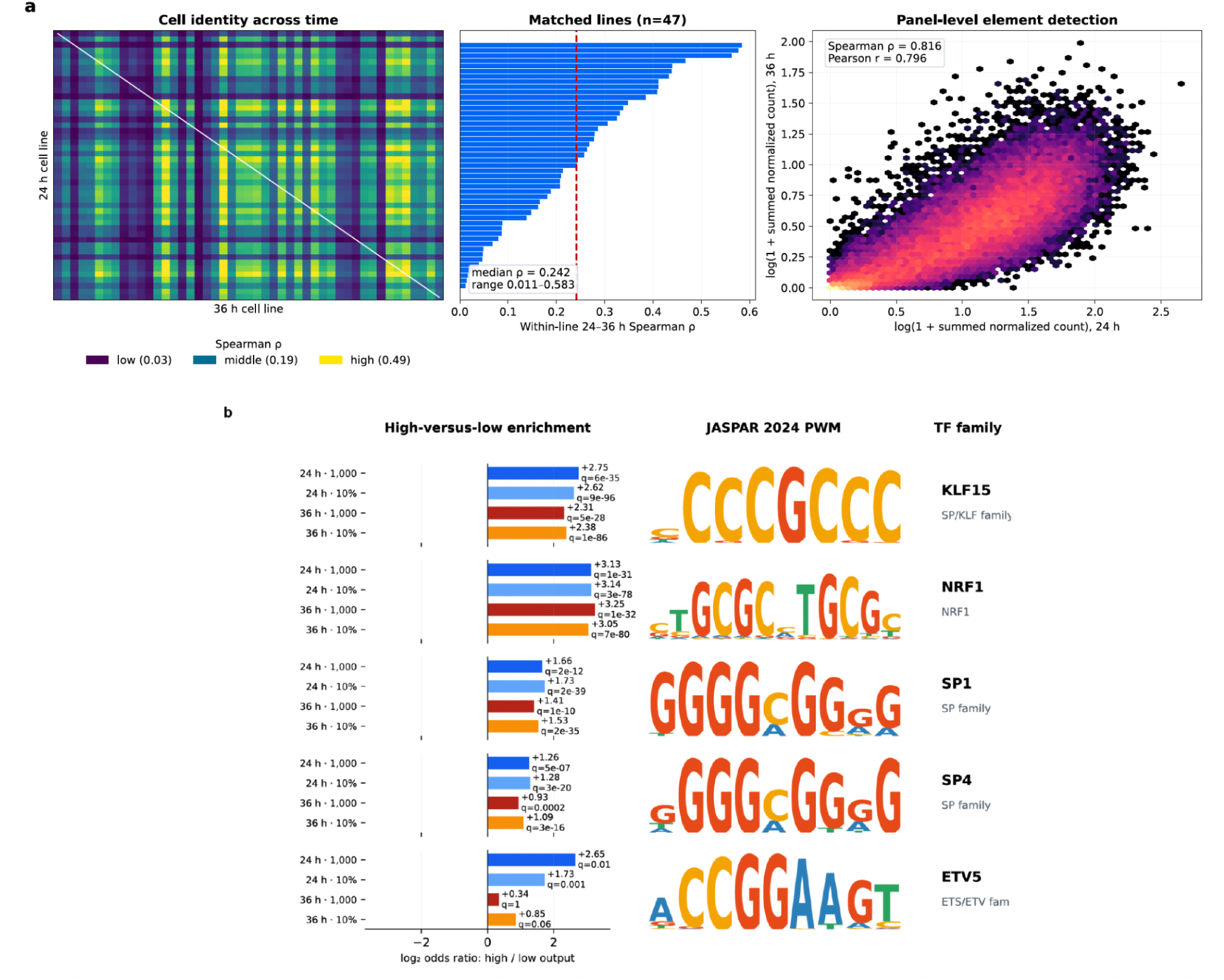
The 5′ UTR/episomal arm is reproducible across time points and enriched for transcription-factor motifs. **a**, Reproducibility between the 24 h and 36 h time points. Left, Spearman correlation between every 24 h and every 36 h cell line, with matched lines located on the diagonal. Middle, within-line 24-36 h correlation across all 47 matched cell lines, ranked, with the median marked (median ρ = 0.242; range, 0.011–0.583). Right, panel-level element detection at 36 h versus 24 h, shown as log(1 + summed normalized count) per element (Spearman ρ = 0.816; Pearson r = 0.796). **b**, Transcription-factor motif enrichment among high-output 5′ UTR elements. Bars give the log_2_ odds ratio for motif occurrence in high-versus low-output elements at two selection thresholds (top/bottom 1,000 elements and top/bottom 10%) and both time points, with q values annotated. JASPAR 2024 position weight matrices and TF family are shown at right. KLF15, NRF1, SP1, and SP4 were consistently enriched across thresholds and time points; ETV5 enrichment was time-dependent and was not significant at either 36 h threshold.

We next asked whether high-output 5′ UTR elements were enriched for known transcription-factor binding motifs. Comparing high- and low-output elements using both the top/bottom 1,000 and top/bottom 10% thresholds identified reproducible enrichment for motifs matching KLF15, NRF1, SP1, and SP4. ETV5 enrichment was time-dependent and not significant at either 36 h threshold (JASPAR 2024 PWMs; **Fig. 4b**). These motifs are largely consistent with earlier MPRA studies of promoter-proximal activity, including lentiMPRA^26^, despite differences in construct design between the two platforms.

### Chronos reveals context-specific *cis*-*trans* regulatory interactions

The resulting Chronos datasets provide compact *cis*-regulatory measurements across matched cell contexts. Each row corresponds to a synthetic regulatory element, each column corresponds to a cell-line context, and each entry represents a final regulatory output, such as DNA-normalized expression or RNA decay rate. These measurements directly represent the interaction between compact *cis*-regulatory sequence and the *trans*-regulatory environment of each cell line.

Across Tahoe’s Mosaic pool, compact regulatory elements exhibited both shared and context-specific behavior (**Figs. 5**, **6**). Some elements were consistently active or stable across many cell lines, whereas others showed stronger effects in specific contexts, as illustrated by representative examples from each library (**Figs. 7**, **8**). This structure is the central signal Chronos is designed to expose: the same compact *cis* element can have different quantitative effects depending on the cellular environment in which it is interpreted.

**Fig. 5.**
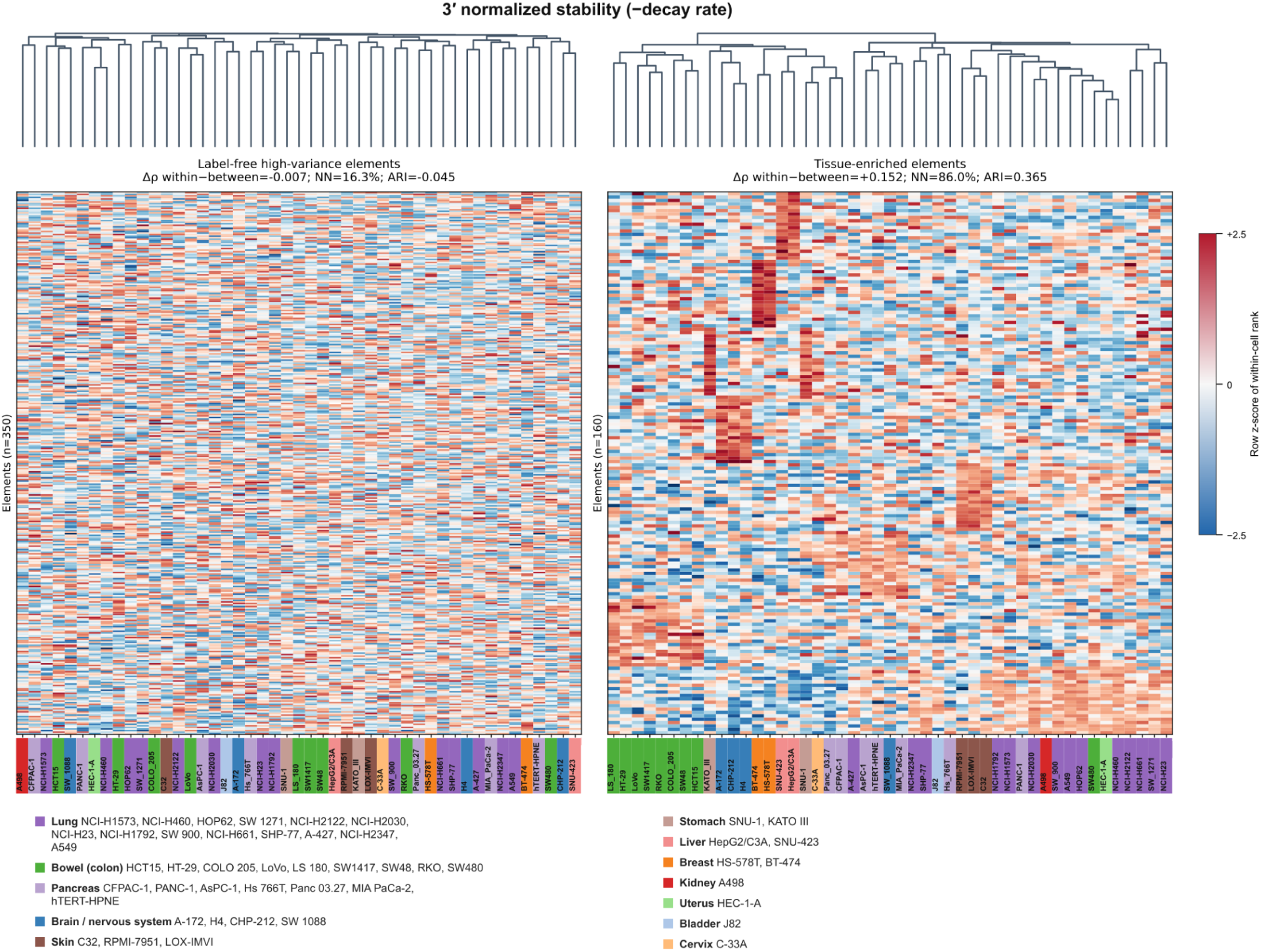
Tissue-enriched 3′ UTR elements recover tissue structure across the cell-line panel without using tissue labels. Hierarchical clustering of 47 cell lines on normalized RNA stability (−k). Left, 350 label-free high-variance elements show no tissue organization (Δρ within − between = −0.007; nearest-neighbor tissue concordance 16.3%; ARI = 0.045). Right, tissue-enriched elements (n = 160) recover tissue structure (Δρ = +0.152; nearest-neighbor concordance 86.0%; ARI = 0.365). Color, row-scaled stability. Column color bars denote tissue of origin.

**Fig. 6.**
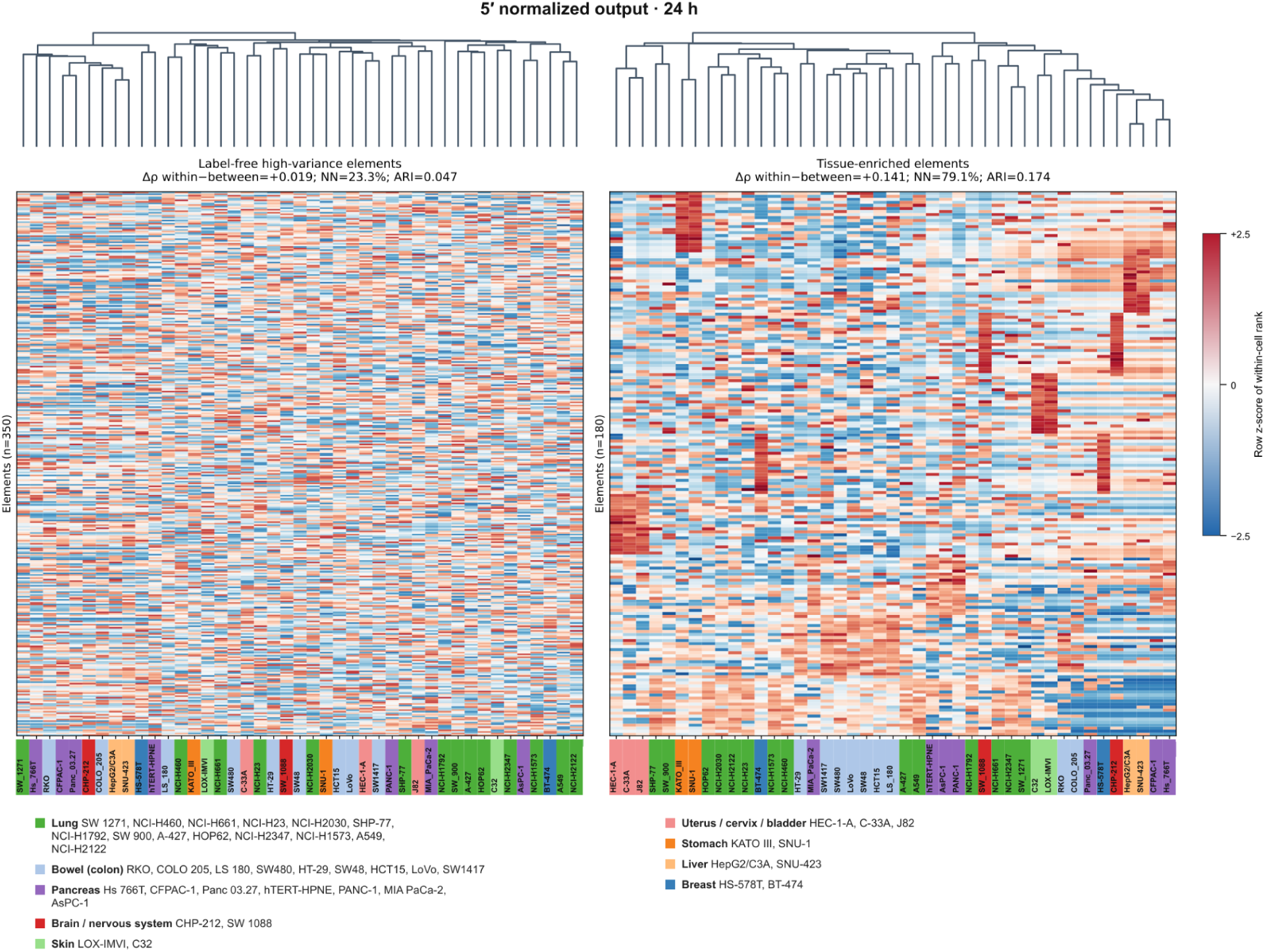
Tissue-enriched 5′ UTR elements recover tissue structure across the cell-line panel. Hierarchical clustering of cell lines on DNA-normalized 5′ UTR output at 24 h. Left, label-free high-variance elements (n = 350; Δρ = +0.019; nearest-neighbor concordance 23.3%; ARI = 0.047). Right, tissue-enriched elements (n = 90; Δρ = +0.141; nearest-neighbor concordance 79.1%; ARI = 0.174). Column color bars denote tissue of origin.

**Fig. 7.**
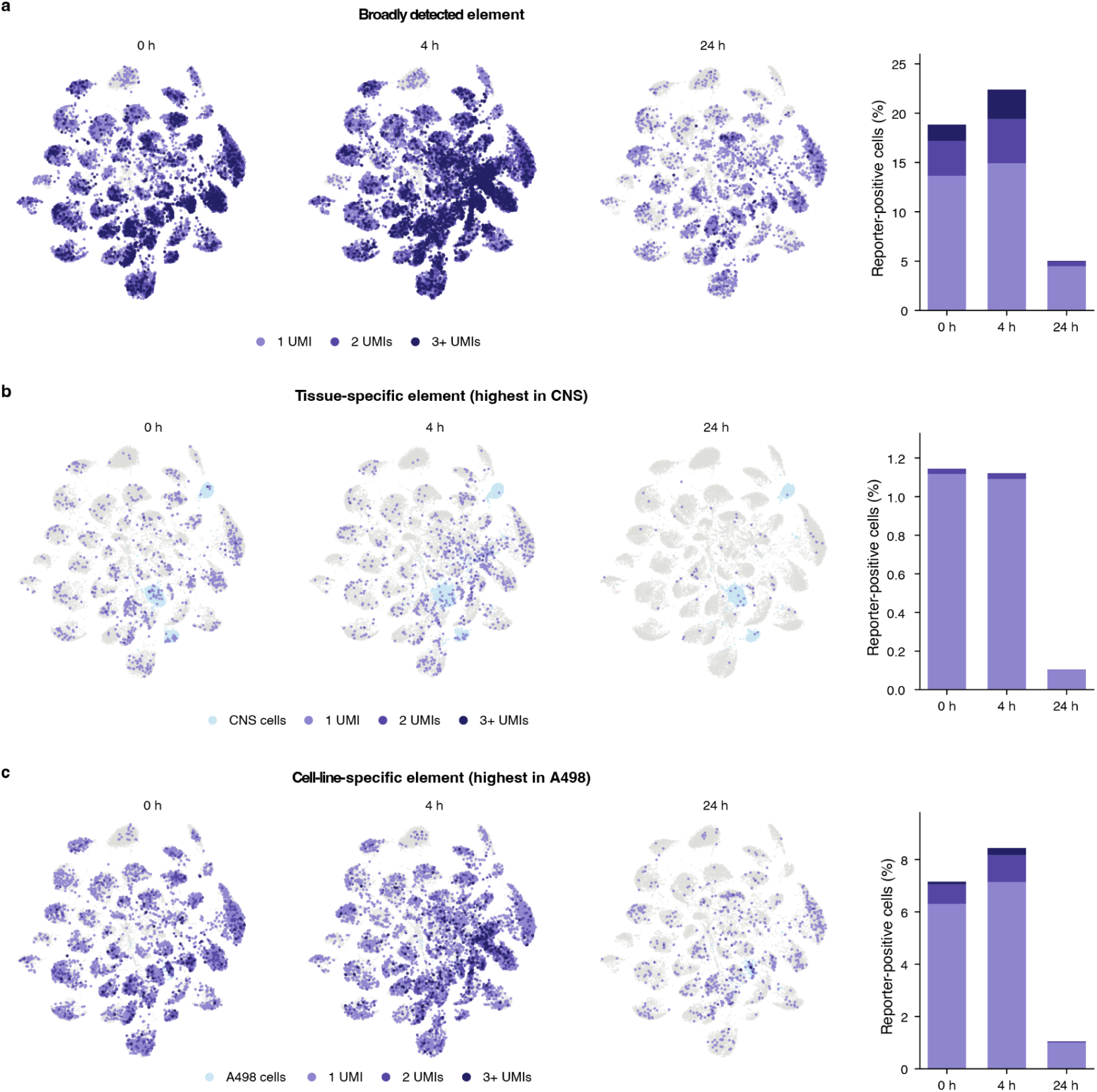
Single 3′ UTR elements range from broadly detected to cell-line-restricted. Representative library members shown on the pooled single-cell UMAP at 0, 4, and 24 h after RNA delivery, with cells colored by synthetic-mRNA UMI count (1, 2, 3+), and the percentage of reporter-positive cells at each time point at right. **a**, A broadly detected element. **b**, A tissue-restricted element. **c**, An element highest in A498. Detection declines by 24 h in all three cases, consistent with decay of the delivered mRNA.

**Fig. 8.**
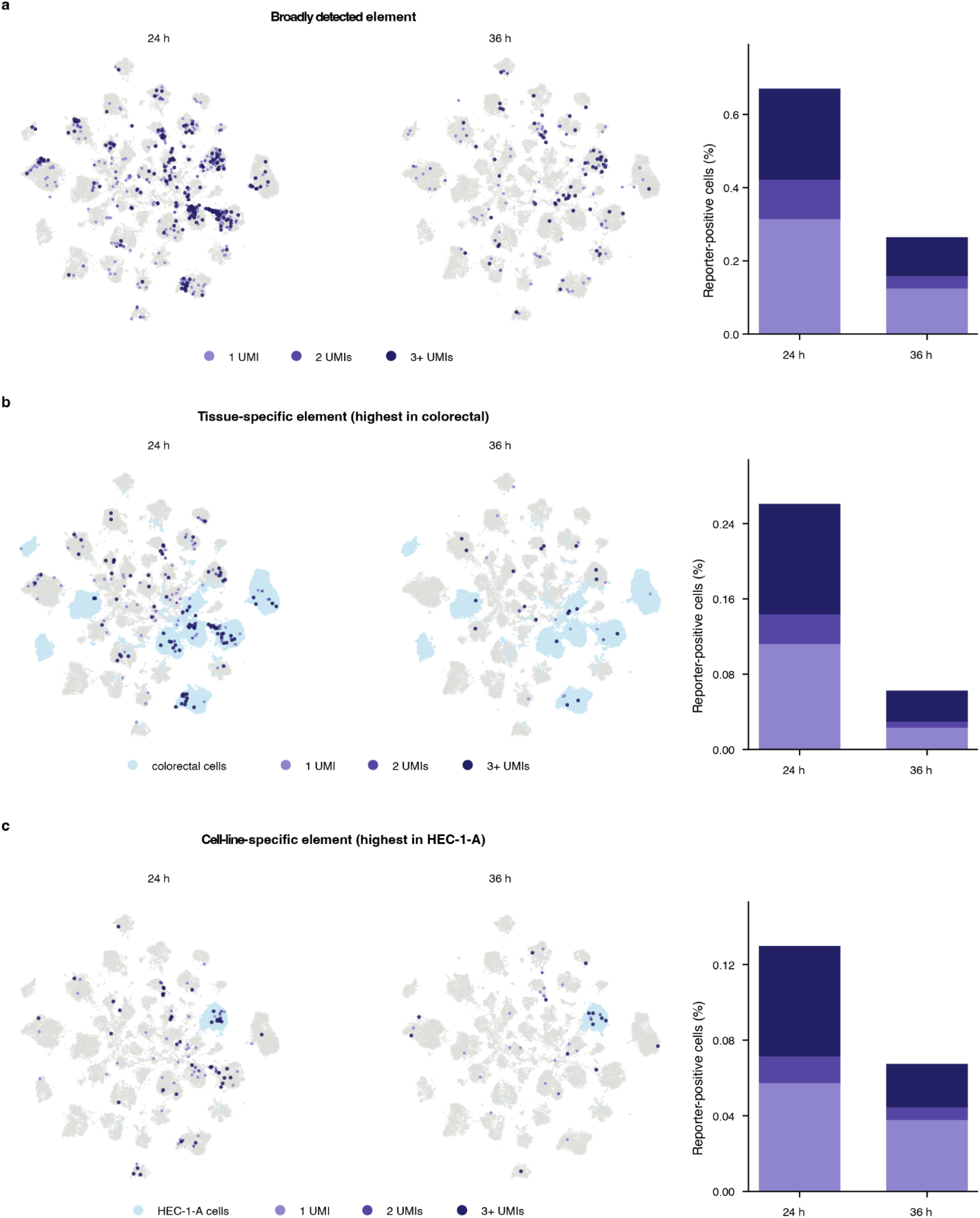
Individual 5′ UTR elements show a comparable range of context restriction. Representative library members on the pooled single-cell UMAP at 24 and 36 h after episomal plasmid delivery. Cells are colored by synthetic-transcript UMI count (1, 2, 3+); in **b** and **c**, cells from the enriched tissue or line are shaded light blue. Bar charts at right show the percentage of reporter-positive cells at each time point, stacked by UMI count. **a**, A broadly detected element. **b**, A tissue-specific element, highest in colorectal lines. **c**, A cell-line-specific element, highest in HEC-1-A.

These context-specific elements support several modeling tasks. Models can learn sequence-to-expression relationships, sequence-to-stability relationships, transfer across cell contexts, and predict where a compact *cis* element will be interpreted differently as the *trans*-regulatory environment changes. Because Chronos expands the measured gene space with defined synthetic genes, these tasks are less confounded by the broad endogenous regulatory architecture that surrounds native genes.

### Chronos reveals sequence motifs underlying both transcriptional and post-transcriptional regulatory control

*De novo* motif discovery across the *Tria-47×28K* decay-rate matrix identified sequence features linked to RNA stability: GC content and A-rich motifs^27,28^ were associated with slower decay, whereas AU content, poly-U tracts, AU-rich elements^29^, and Pumilio-like motifs^30^ were associated with faster decay (**Fig. 9a**). Among *de novo* motifs lacking confident RBP-PWM matches (**Fig. 9b**), two were compatible with established miRNA-response elements. TACCTCA corresponds to a let-7-family-compatible site^31^, whereas TGTAGCA corresponds to a miR-221-family-compatible site^32^. Their enrichment among faster-decaying elements is consistent with miRNA-mediated RNA destabilization. TGCTGCT lacked a confident RBP or canonical miRNA-site assignment and was therefore retained as an unassigned decay-associated *cis*-regulatory motif. *De novo* motif discovery independently recovered the same axis across nearly all cell-line contexts, indicating that Chronos captures a reproducible *cis*-regulatory grammar for RNA stability.

**Fig. 9.**
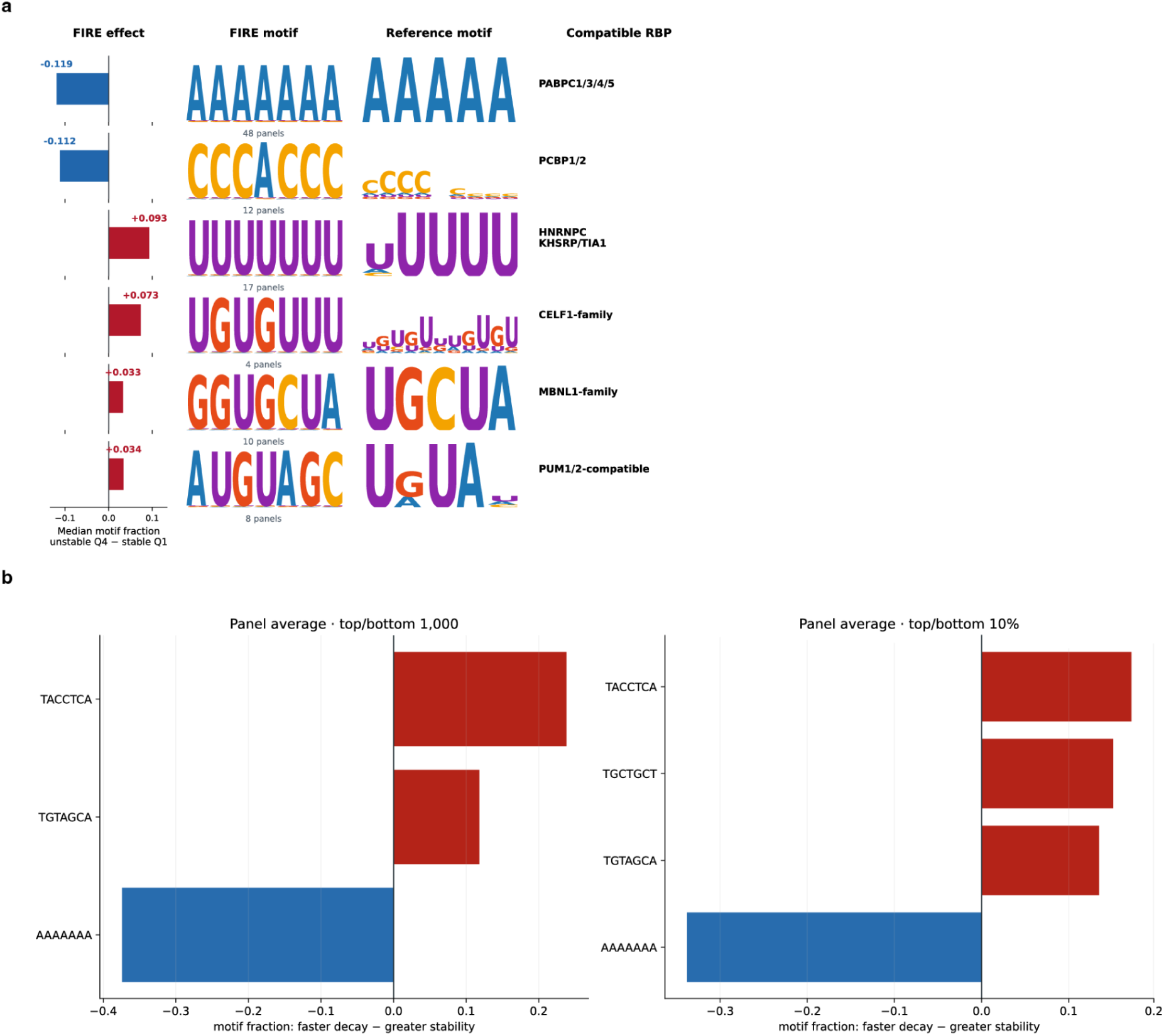
*De novo* motif discovery recovers a coherent stability grammar in 3′ UTRs. **a**, Motifs discovered *de novo* by FIRE across cell-line panels. For each motif, the bar gives its FIRE effect: the median difference in motif-containing fraction between the least stable (Q4) and most stable (Q1) decay quartiles, so negative values indicate enrichment among stable elements. The discovered motif, its closest reference motif, and its compatible RNA-binding proteins are shown at right. The number beneath each logo is how many cell-line panels recovered that motif. Two motifs are enriched among stable elements: an A-rich motif matching PABPC1/3/4/5 and a C-rich motif matching PCBP1/2. Four are enriched among unstable elements: a poly-U motif matching HNRNPC and KHSRP/TIA1, two UG-rich motifs matching the CELF1 and MBNL1 families, and a UGUA-containing motif compatible with PUM1/2. **b**, Panel-average motif fraction in faster-decaying minus more stable elements, at two selection thresholds: the top and bottom 1,000 elements on the left, and the top and bottom 10% on the right. TACCTCA and TGTAGCA are associated with faster decay at both thresholds; TGCTGCT is associated with faster decay only at the 10% threshold. AAAAAAA is strongly associated with greater stability.

Motivated by the independent recovery of let-7- and miR-221/222-compatible motifs from unbiased *de novo* discovery, we next asked whether canonical seed-matched sites for these two families showed a systematic relationship with decay across the full panel. Elements carrying canonical let-7^31^ and miR-221/222^32^ seed-matched sites, classified as 8mer, 7mer-m8, or 7mer-A1 sites as applicable^33^, decayed faster (**Fig. 10a**). To test whether this relationship extended beyond these two families, we then computed a global miRNA site burden per element, summing predicted seed-matched sites across all annotated human miRNAs and weighting each site by the endogenous expression of its cognate miRNA in the corresponding cell-line context. This context-weighted global burden showed a consistent positive association with decay rate across the panel (**Fig. 10b**).

**Fig. 10.**
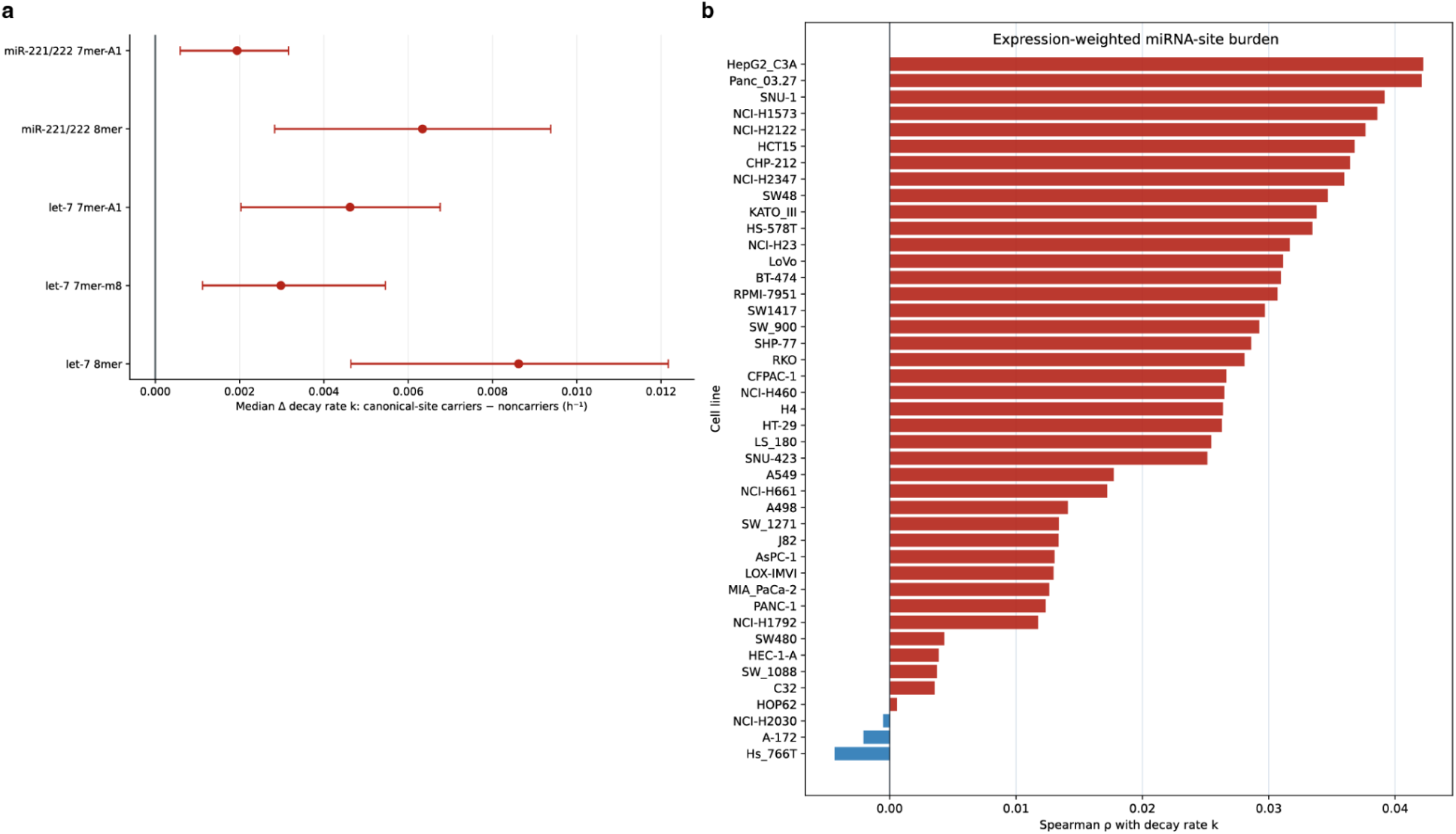
Canonical miRNA target-site effects and expression-weighted site burden across cell lines. **a**, Median difference in decay rate between carriers and non-carriers of canonical let-7 and miR-221/222 sites, by site class (8mer, 7mer-m8, 7mer-A1); points are medians, bars are confidence intervals. All five classes shift toward faster decay. **b**, Spearman correlation between expression-weighted miRNA-site burden and decay rate k, per cell line. The association is positive in the large majority of contexts.

Beyond miRNA-mediated regulation, we asked whether the same decay-rate matrix could be explained by RNA-binding protein activity: these sequence features also aligned with known *trans*-regulatory programs annotated using 544 CISBP-RNA PWMs^34^, with elements enriched for sequences compatible with annotated stabilizing RBPs showing slower decay, while those compatible with annotated destabilizing RBPs trended toward faster decay (**Fig. 11**).

**Fig. 11.**
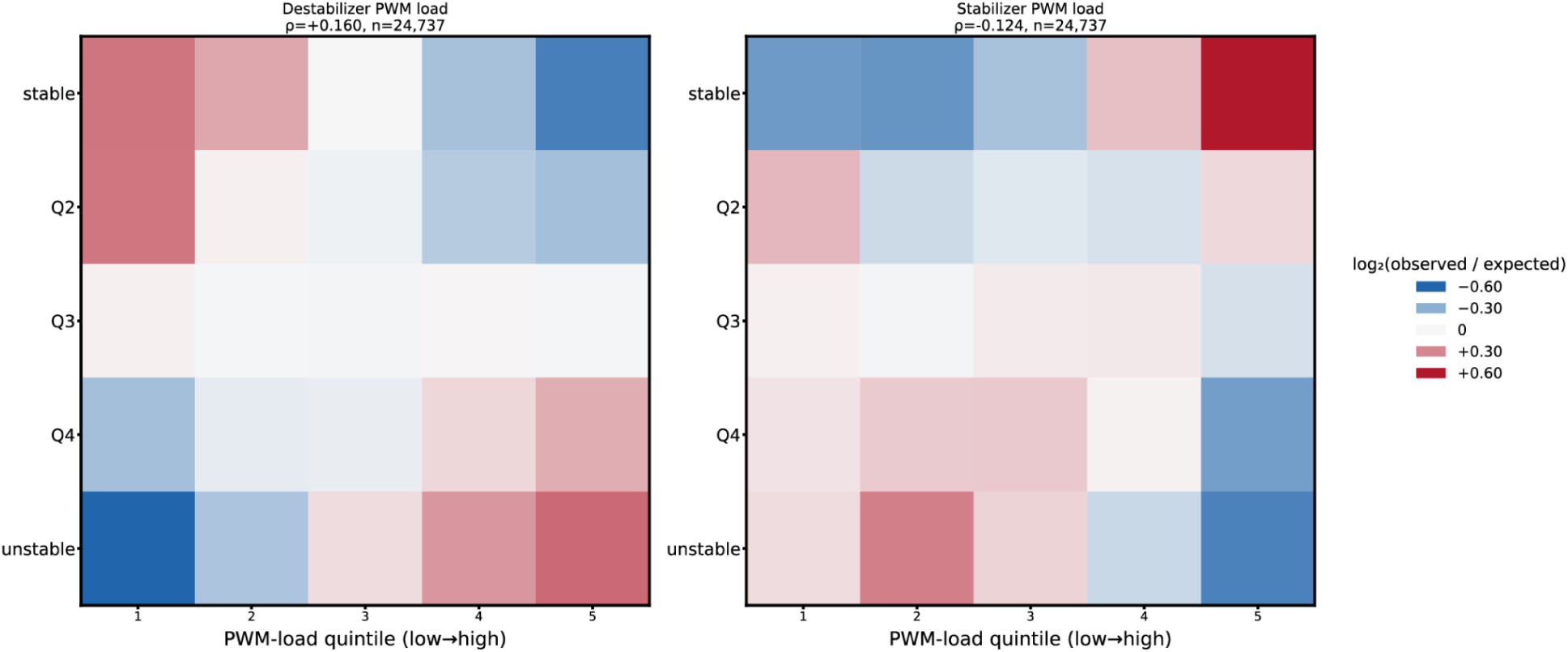
Stabilizing and destabilizing RBP motif loads shift decay in opposite directions across the panel. Decay-rate quartile (stable to unstable) against PWM motif-load quintile (low to high), pooled across cell lines, for annotated destabilizing (left, +0.160) and stabilizing (right, −0.124) RNA-binding proteins; n = 24,737 elements. Color, log_2_(observed/expected).

Applying the same *de novo* motif discovery approach to the 5′ UTR/episomal library recovered a distinct signature: GC-rich motifs enriched among the highest-expressing elements (**Fig. 12a, b**), consistent with the transcription-factor motif enrichment identified independently above (**Fig. 4b**). Sparse-autoencoder features derived from Evo 2 layer-26 representations converged on these same GC-rich regulatory programs, providing an orthogonal, motif-agnostic confirmation of this signal (**Fig. 12c**). GC/CpG-rich sequences can influence gene output through both promoter-driven transcriptional activity and mRNA stability^35^, and because the episomal/plasmid format captures both layers jointly, this dual role likely contributes to the GC-rich signal recovered here.

**Fig. 12.**
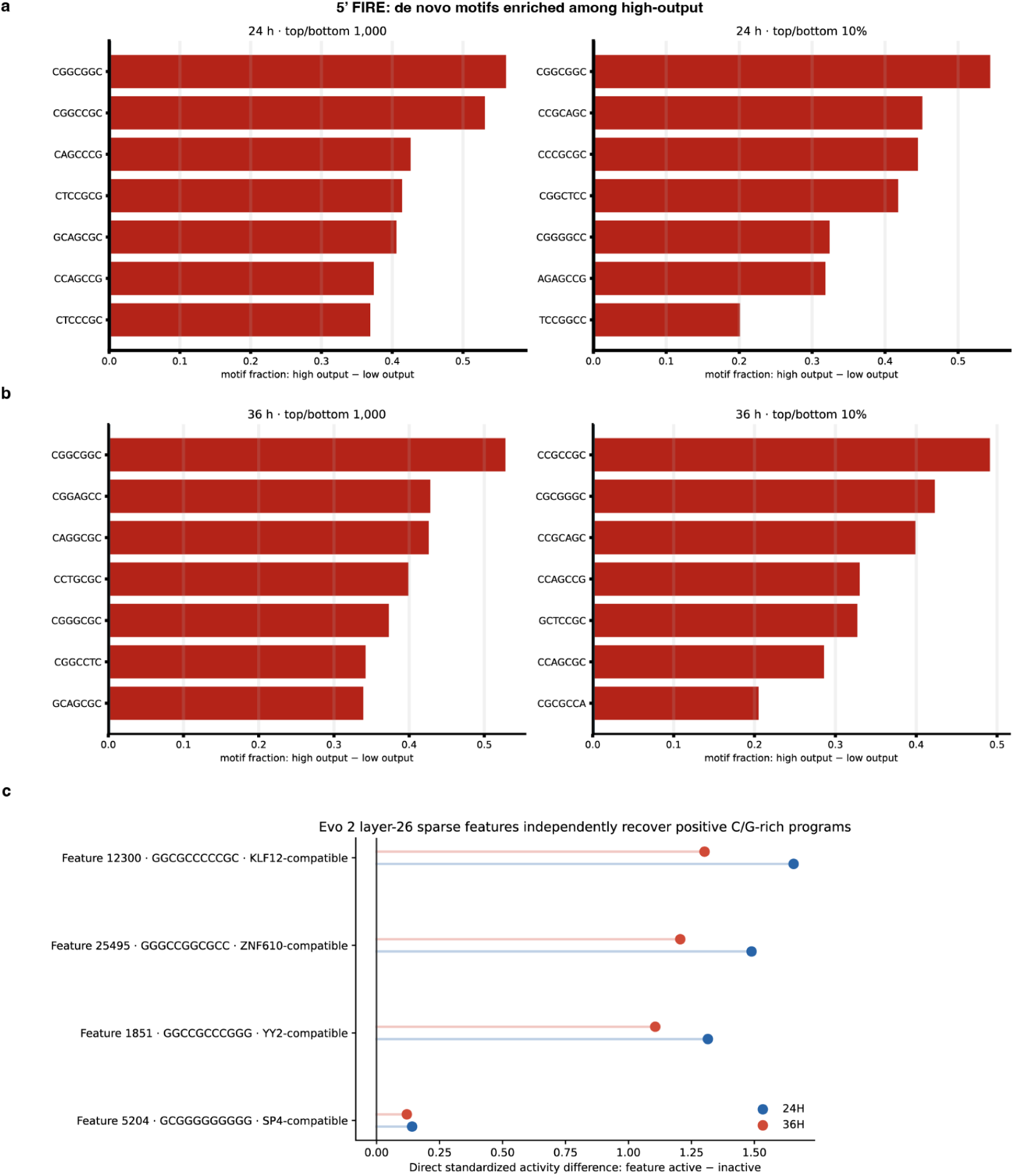
High-output 5′ UTR elements converge on GC/CpG-rich sequence by two independent routes. **a,b**, Motifs discovered *de novo* by FIRE among high-output elements, at 24 h in a and 36 h in b. Each panel uses two selection thresholds: the top and bottom 1,000 elements on the left and the top and bottom 10% on the right. Bars give the motif fraction in high-output minus low-output elements. Every motif recovered is C/G-rich. **c**, Sparse features from layer 26 of an internal Evo 2 model recover the same programs without motif input. For each feature, points give the standardized difference in activity between elements where the feature is active and elements where it is inactive, at 24 h and 36 h. Three of the four features are C/G-rich and compatible with KLF12, ZNF610, and YY2. The fourth, an SP4-compatible poly-G feature, shows a much smaller effect.

### Chronos provides a compact *cis*-regulatory counterpart to perturbation atlases and virtual-cell models

Large perturbation atlases measure how endogenous transcriptomes respond to changes in *trans*-regulatory state. *Tahoe-100M*^1^ is one example, focused on chemical perturbations across a large cancer cell-line panel, but the same conceptual limitation applies to genetic perturbation atlases and to models trained on them. STATE^10^, STACK^11^, Geneformer^8^, scGPT^9^, scFoundation^36^, and related models can learn how cellular state changes after perturbation or across biological contexts, yet their transcriptomic input and output spaces remain anchored to endogenous genes within broad and entangled regulatory contexts.

Chronos adds an auxiliary regulatory lens to this framework. It creates tens of thousands of additional synthetic genes and measures how compact *cis*-regulatory variants are interpreted by each cellular context. A model trained only on endogenous genes must learn sequence regulation from a small and highly confounded set of natural examples. Chronos reduces this problem by holding most of the synthetic gene constant and varying a short regulatory sequence. This does not replace endogenous perturbation atlases; it complements them by adding a controlled sequence-to-function axis. This complementarity creates a natural strategy for joint modeling. Perturbation atlases teach models how endogenous transcriptomes change when the *trans*-regulatory state is altered, whereas Chronos teaches models how controlled changes in *cis*-regulatory sequence are interpreted across cellular contexts. Integrating these datasets would combine the biological breadth of endogenous perturbation measurements with the controlled sequence variation provided by synthetic genes. This creates a progression of increasingly difficult prediction tasks: predicting a new regulatory sequence in a previously observed cellular context; predicting a previously observed sequence in a new cellular context; and, ultimately, predicting a new sequence in a new cellular context. Achieving this final form of compositional generalization would require models to learn both *cis*-regulatory sequence grammar and a transferable representation of *trans*-regulatory cellular state (**Fig. 13**).

**Fig. 13.**
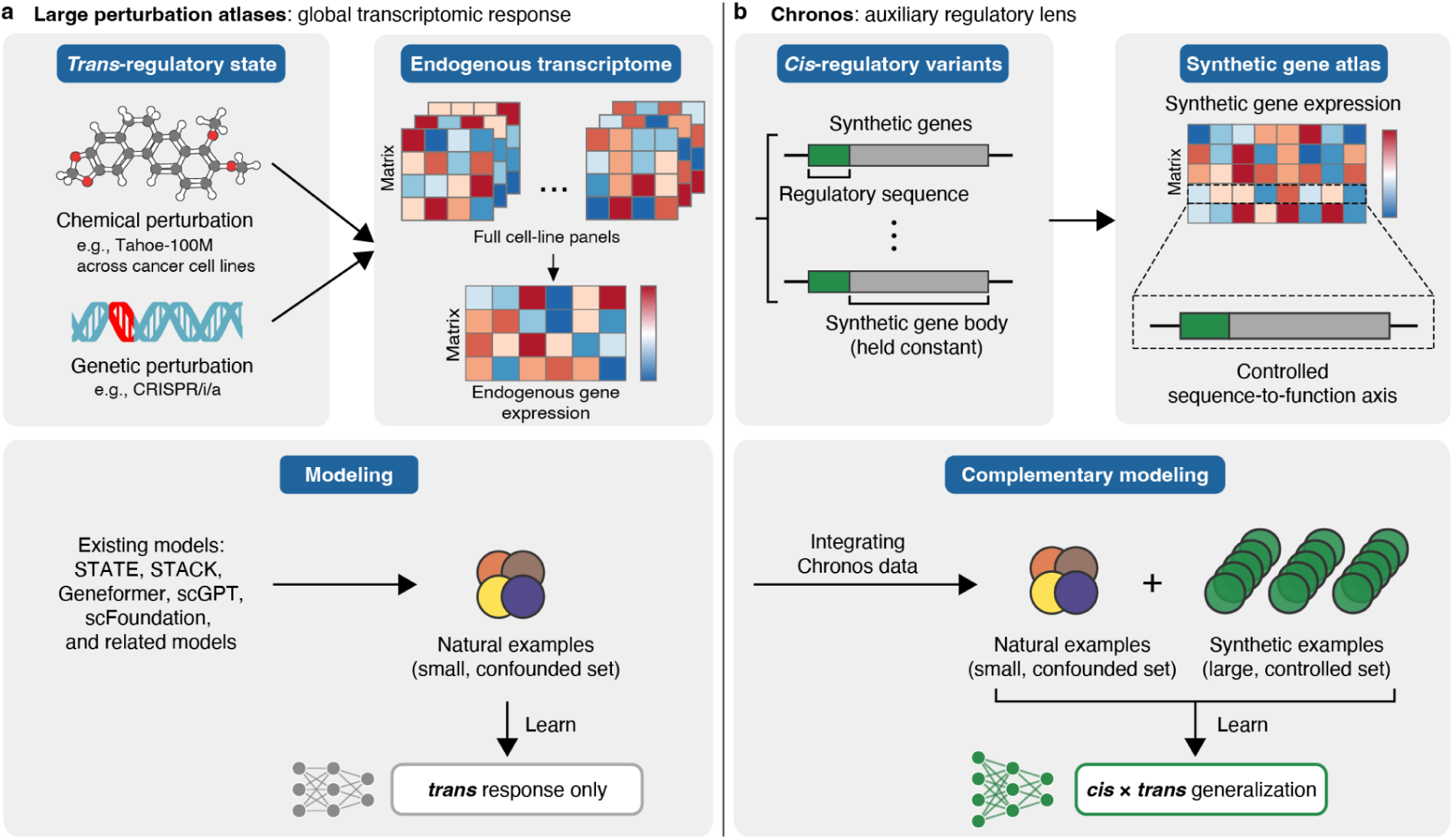
Chronos provides a controlled *cis*-regulatory complement to large perturbation atlases. **a**, Chemical and genetic perturbation atlases measure transcriptome-wide responses to changes in *trans*-regulatory state and provide training data for virtual-cell models. **b**, Chronos holds the synthetic gene architecture largely constant while varying compact regulatory sequences, generating controlled sequence-to-function measurements across cellular contexts. Integrating these complementary datasets could support increasingly difficult prediction tasks: new regulatory sequences in observed cellular contexts, observed sequences in new cellular contexts, and ultimately new sequences in new cellular contexts. This framework aims to combine *cis*-regulatory sequence grammar with transferable representations of *trans*-regulatory cellular state.

Together, *Penta-47×27K* and *Tria-47×28K* provide a matched *cis*-regulatory counterpart to large *trans*-perturbation resources. Direct RNA-delivery experiments (*Tria-47×28K*) report RNA stability and post-transcriptional activity. Episomal/plasmid experiments (*Penta-47×27K*) report DNA-normalized transcriptional output together with RNA-level regulation. In combination, these measurements enable modeling of transcriptional and post-transcriptional programs within the same cellular frame.

### Chronos enables cell-context-conditioned prediction of regulatory activity from UTR sequence

We asked whether a compact UTR sequence alone could predict regulatory activity across the cellular contexts measured by Chronos. Separate models were trained for *Penta-47×27K* and *Tria-47×28K*, which measure normalized expression for 5′ UTR/internal-promoter elements and RNA decay for 3′ UTR elements, respectively. Both models used a compact convolutional architecture derived from PARADE^27^/LegNet^37^, comprising a shared sequence-processing backbone and cell-line-specific ordinal output heads.

Sequences were partitioned into mutually exclusive training, validation, and test sets, with all measurements associated with a sequence assigned to the same partition. Classification thresholds were estimated exclusively from training data. Model selection used validation performance, and the test partitions were held out from model fitting and selection.

For *Penta-47×27K*, 24 h activity was normalized to representation in the input library. Elements with pooled T0 DNA counts of at least 600 were retained, yielding 27,631 unique 50-nt elements. Cell lines with a median positive 24 h pseudobulk RNA count of three or fewer in the training partition were excluded, retaining 29 contexts. NCI-H1792 was subsequently excluded because at least one-third of its training RNA measurements were tied at zero, leaving 28 modeled cell-line contexts. The sequence-level split contained 22,204 training, 2,645 validation, and 2,782 test elements. Within each retained cell line, the 33rd and 67th percentiles of its training measurements defined low-, intermediate-, and high-activity classes. These thresholds were applied unchanged to validation and test measurements.

Five models were trained from independent random initializations for up to 100 epochs with early stopping. Predictions were averaged across the five models using an ensemble policy fixed before test evaluation. A single ensemble predicts cell-line-specific activity class for all 28 retained contexts simultaneously, from sequence alone. On the locked test set, 86.5% of held-out element–cell-line predictions fell within one ordinal class of the measured class, with an ordinal mean absolute error of 0.669. Exact three-class accuracy was 0.467, compared with a nominal three-class chance rate of 0.333, and median per-cell-line macro-F1 was 0.464 (validation accuracy 0.476, median macro-F1 0.471, **Fig. 14**).

**Fig. 14.**
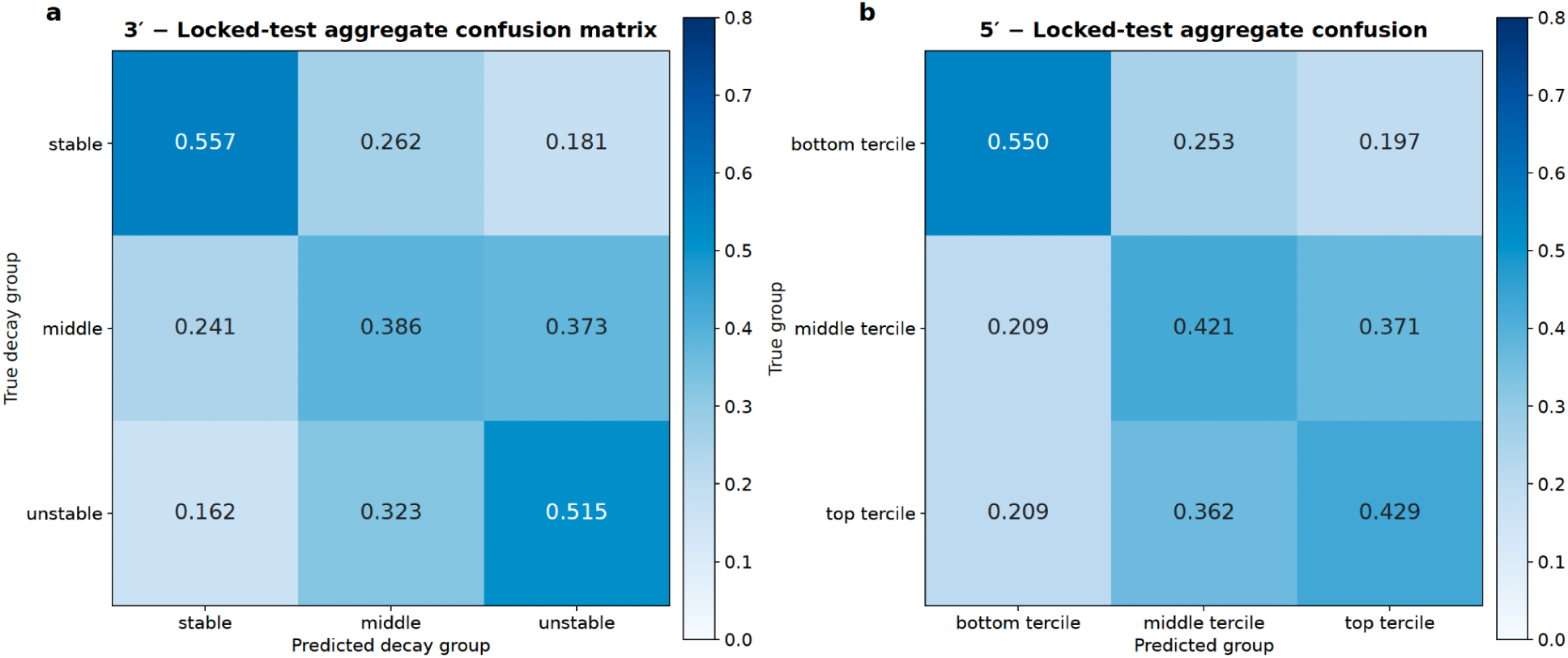
Sequence-only models predict cell-line-specific regulatory class on locked test sets. Row-normalized aggregate confusion matrices for the held-out test partitions, pooled across cell lines. **a**, 3′ UTR stability across 47 cell-line contexts, with elements grouped as stable, middle, or unstable. **b**, 5′ UTR output across 28 contexts, grouped by tercile. In both arms, correct extreme-class assignments were more frequent than assignments to the opposite extreme.

For *Tria-47×28K*, the modeling dataset contained 26,958 unique 240-nt elements, 47 cell lines, and 1,218,278 observed element–cell-line decay rates. The sequence-level split contained 21,621 training, 2,667 validation, and 2,670 test elements. Within each cell line, the 33rd and 67th percentiles of its finite training decay measurements defined stable, intermediate, and unstable classes. Missing measurements were masked during training and evaluation.

The 3′ UTR model used sequence as its only input and produced two ordinal logits for each cell line. Five independently initialized models were trained and combined using the same prespecified mean-ensemble policy. As in the 5′ arm, a single ensemble predicts cell-line-specific stability class for all 47 contexts simultaneously, from sequence alone. On the locked test set, 88.54% of held-out element–cell-line predictions fell within one ordinal class of the measured class, with an ordinal mean absolute error of 0.628. Exact three-class accuracy was 0.486, against a chance rate of 0.333, and median per-cell-line macro-F1 was 0.481 (validation accuracy 0.478, median macro-F1 0.478, **Fig. 14**).

## Discussion

Chronos addresses a central gap in current virtual-cell datasets and models. Large perturbation atlases measure how cells respond to changes in *trans*-regulatory state, but they do not densely sample the *cis*-regulatory sequences that cells must interpret. Models trained on these resources inherit the same constraint: the measured gene space is limited to endogenous genes, and each endogenous gene carries a broad regulatory context that is difficult to disentangle. Chronos provides a complementary strategy by generating synthetic genes with compact variable regulatory windows and measuring their behavior across many matched cellular contexts. Internally, Chronos is being extended toward more clinically relevant, therapeutically oriented systems (e.g., PBMCs, organoids, or *in vivo* models) using LNP-formulated mRNA delivery, capabilities that fall outside the scope of the public release.

The generation of *Penta-47×27K* and *Tria-47×28K* demonstrates the feasibility of this approach in a pooled cancer cell-line system, testing whether this region can regulate transcription in addition to its established role in translation. In one pooled experiment, Chronos measures compact regulatory elements across approximately 50 cellular contexts, using whole-transcriptome profiles to assign context and targeted synthetic-gene reads to quantify regulatory output. In the RNA-stability arm (*Tria-47×28K*), Chronos estimates decay rates from longitudinal RNA-delivery measurements. In the transcriptional arm (*Penta-47×27K*), Chronos estimates DNA-normalized expression from episomal/plasmid delivery, testing whether the 5′ UTR can regulate transcription in addition to its established role in translation. The resulting dataset supports direct modeling of how short regulatory sequences interact with cellular context to determine gene expression output.

Several aspects of the release are important for interpretation. First, Chronos uses single-cell profiling to identify cell context, but the synthetic regulatory output is a context-level measurement. Each cell contains only part of the synthetic library, so the relevant unit of analysis is the collapsed cell-line or cell-type stratum rather than an individual cell. Second, the data release intentionally focuses on final, modeling-ready regulatory outputs: decay rates and DNA-normalized expression estimates. Third, the RNA-delivery datasets measure synthetic mRNA behavior after delivery of chemically defined RNA, specifically mRNA incorporating N1-methylpseudouridine, a modification widely used in mRNA therapeutics. Because this is the same chemically modified mRNA used in the clinic, these measurements directly reflect the regulatory landscape therapeutic developers actually need to model, including emerging applications such as *in vivo* CAR-T cell engineering^38^. They may, however, differ from endogenous unmodified mRNA regulation, a distinction also recognized by bulk methods such as PERSIST-seq^39^ and viral-derived RNA screens^40^ that similarly incorporate modified mRNA into their platforms.

By releasing Therna Biosciences’ Chronos platform within a cellular frame that integrates with existing perturbation resources, we aim to make *cis*-regulatory modeling directly usable by the virtual-cell community. The dataset supports sequence-to-function learning, cell-context-conditioned prediction, transfer across cell lines, and benchmarking of models that claim to represent regulatory biology. More broadly, Chronos illustrates a scalable experimental principle: when the natural regulatory genome is too large and too sparsely sampled, synthetic genes can expand the measured gene space while compressing the variable *cis* code into a form that can be measured, learned, and integrated with cell-state atlases across dozens of contexts at once.

## Data availability

The Chronos datasets are publicly available through the Therna Hugging Face collection (https://huggingface.co/collections/therna/chronos). The accompanying modeling notebooks, including an environment for reproducing the released analyses, are available through the Chronos public release template on Lightning AI (https://lightning.ai/thernabiosciences/templates/chronos-public-release).

## Acknowledgments

We gratefully acknowledge the Tahoe team for enabling this study by providing access to the Mosaic cancer cell-line pool used in *Tahoe-100M* and their continued support of open projects. We also thank the Tahoe team for sharing and supporting the processing and cell-line assignment workflow used to map Chronos whole-transcriptome profiles back to their matched cell-line contexts, including the *Tahoe-100M*-derived reference data and scANVI-based assignment pipeline. This support enabled evaluation of Therna Biosciences’ Chronos platform across approximately 50 cellular contexts in a single pooled experiment and alignment of the resulting *cis*-regulatory measurements with the broader Tahoe perturbation resource. We thank Parse Biosciences and 10x Genomics for their support of this project. HG is an Arc Core Investigator, but this work was entirely funded by Therna Biosciences.

## Extended Data Figures

**Extended Data Fig. 1.**
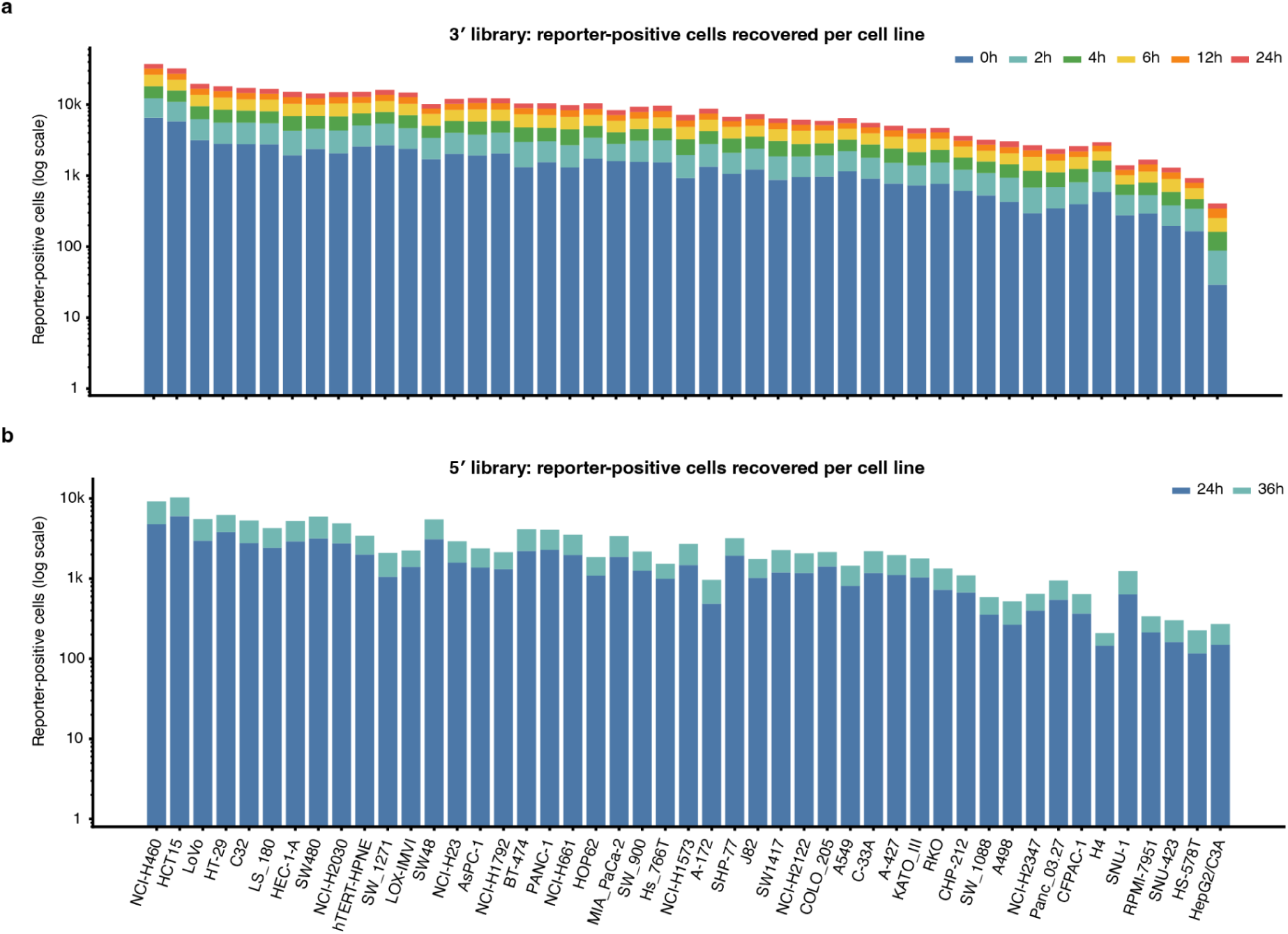
Reporter-positive cell recovery across the pooled cell-line panel over time in both delivery formats. **a,** Number of reporter-positive cells recovered for each annotated cell line in the 3′ UTR library at 0, 2, 4, 6, 12, and 24 h. **b,** Number of reporter-positive cells recovered for each annotated cell line in the 5′ UTR library at 24 and 36 h. In both panels, bars are stacked by time point, ordered by total recovery, and displayed on a logarithmic scale. Differences reflect the combined effects of delivery, cell recovery, sequencing depth, and reporter detection and should not be interpreted as calibrated transfection efficiencies.

## Methods

### Synthetic gene library design

*Penta-47×27K* and *Tria-47×28K* include compact synthetic gene libraries designed to measure transcriptional and post-transcriptional regulatory output. The RNA-stability library (*Tria-47×28K*) includes synthetic mRNA variants with constant coding and adapter regions and compact variable UTR regions followed by a poly(A) tail. The transcriptional library (*Penta-47×27K*) includes compact *cis*-regulatory fragments in episomal/plasmid format, with fragment-specific DNA representation measured to support DNA-normalized expression estimates. In the DNA-delivery format, measured synthetic-RNA abundance reflects transcriptional activity together with post-transcriptional regulation, whereas RNA-delivery experiments more directly measure RNA stability and post-transcriptional effects. Comparing RNA and DNA delivery formats allows compact sequence effects to be decomposed across regulatory layers.

Specific implementations include a 3′ UTR RNA-stability library comprising approximately 30,000 synthetic mRNA variants with a constant 50 nt natural *HBB* 5′ UTR sequence and variable 3′ UTR regions of approximately 240 nt. Prior to treating the cells, seven control RNAs with unique 8 nt barcodes in their 3′ UTRs were spiked into the library at varying concentrations: 4-, 2-, 1-, 0.5-, 0.25-, 0.125-, and 0.0625-fold of the average library member abundance. As stability controls, the library also included two 3′ UTR control elements that stabilize mRNA along with destabilizing point mutants. The release also includes a 5′ UTR/internal-promoter library comprising approximately 30,000 compact synthetic gene variants containing approximately 50 nt variable regions. In the episomal/plasmid design, fragment-specific RNA counts are normalized to fragment-specific DNA concentration in the pool to estimate expression output per input DNA molecule.

### Synthetic RNA delivery

Synthetic RNAs were generated with CleanCap Reagent AG (Trilink Biotechnologies, Cat. No. N-7113) and N1-methylpseudouridine. The Tahoe Mosaic cell-line pool provided by the Tahoe team was cultured in RPMI 1640 Medium (Gibco, Cat. No. 11875093) supplemented with 10% FBS, 100 U/mL penicillin, and 100 μg/mL streptomycin under standard tissue culture conditions (37°C, 5% CO_2_). Cells were seeded in 12-well plates at a density of approximately 400,000 cells per well and treated with the synthetic RNA library using Lipofectamine MessengerMAX (Invitrogen, Cat. No. LMRNA015). Samples were harvested in biological duplicate as untreated controls and at 0, 2, 4, 6, 12, and 24 hours post-wash using TrypLE Express Enzyme (Gibco, Cat. No. 12604013). For all treated conditions, the transfection mixture was removed after a 4-hour incubation period, cells were washed twice with D-PBS (Gibco, Cat. No. 14190094), and fresh complete media was added.

### Episomal and plasmid delivery

Compact *cis*-regulatory libraries, cloned downstream of an EF1α core promoter region, were delivered as episomal or plasmid DNA. Cells were seeded in a 6-well plate format at a density of approximately 880,000 cells per well and transfected in biological duplicate using FuGENE 4K Transfection Reagent (Promega, Cat. No. E5911). Following a 5-hour incubation, cells were washed twice with PBS and supplied with fresh culture medium. At the indicated post-wash time points (24 h or 36 h), cells were harvested using TrypLE Express Enzyme (Gibco, Cat. No. 12604013) and immediately prepared for single-cell library construction using the 10x Genomics workflow.

### Target-specific digital PCR quantification (RNA delivery)

Total RNA was extracted using the PureLink RNA Mini Kit (Invitrogen, Cat. No. 12183025) with DNase I digestion. Reverse transcription was performed using SuperScript IV Reverse Transcriptase (Invitrogen, 18090200) primed with a 1:1 mixture of gene-specific primers targeting human GAPDH and the synthetic mRNA library. Following cDNA synthesis, RNase H (Invitrogen, Cat. No. 18021071) was added to digest the RNA template within RNA:DNA hybrids. Absolute target concentrations were quantified by digital PCR on a QIAcuity instrument using QIAcuity 3X EvaGreen PCR Master Mix (Qiagen, Cat. No. 250112). Two separate reactions were performed per sample using primer pairs specific to either GAPDH or the synthetic mRNA library, and final absolute concentrations (copies/µL) were calculated with QIAcuity Software using Poisson modeling.

### Single-cell library preparation and sequencing (RNA delivery)

Single-cell libraries were prepared using CRISPR Detect for the Parse Evercode WT Mega workflow (version 3) with custom modifications for the synthetic RNA library. After Barcoding Round 3, a total of 885,000 fixed cells were recovered and used as input for the 3′ UTR experiment. Fifteen sublibraries were generated, each with approximately 59,000 cells.

Synthetic mRNA library members were enriched to create 15 indexed enrichment libraries, and standard whole-transcriptome libraries were generated for endogenous transcriptome analysis.

Sequencing libraries were pooled, aiming for a target sequencing depth of 15,000 reads per cell for the whole transcriptome sublibraries and 10,000 reads per cell for the enrichment sublibraries. Sequencing was performed on a NovaSeq X Plus 25B flow cell using a 300-cycle kit in paired-end format with 5% PhiX. The run used 264 cycles for Read 1, 8 cycles for Index 1, 8 cycles for Index 2, and 58 cycles for Read 2.

### Single-cell library preparation and sequencing (plasmid delivery)

Single-cell libraries were prepared using 10x Chromium GEM-X Single Cell 5’ Reagent Kits, including Feature Barcode technology with custom modifications for the synthetic library. Two microfluidics chips were loaded with cell samples added to six wells each on a Chromium iX machine, aiming for a total of 240,000 single cells. Sequencing libraries were pooled, aiming for a target sequencing depth of 30,000 reads per cell for the standard gene expression library and 20,000 reads per cell for the enrichment library. Sequencing was performed on four lanes of a NovaSeq X Plus 25B flow cell using a 100-cycle kit in paired-end format with 1% PhiX. The run used 28 cycles for Read 1, 10 cycles for Index 1, 10 cycles for Index 2, and 90 cycles for Read

To establish a baseline DNA delivery, cells were washed and harvested immediately following a 5-hour plasmid DNA transfection. Plasmid DNA was extracted using the Quick-DNA MiniPrep Kit (Zymo Research, Cat. No. D3025). Extracted plasmid DNA samples and the input plasmid library were prepared for next-generation sequencing via PCR amplification using custom primers and NEBNext Ultra II Q5 Master Mix (New England Biolabs, Cat. No. M0544X). Sequencing was performed on an Illumina NovaSeq X Plus system using a 10B flow cell with a 100-cycle kit in single-end format, spiked with 10% PhiX control. Run cycle parameters were set to 101 cycles for Read 1, 12 cycles for Index 1, and 24 cycles for Index 2.

### Whole transcriptome processing and cell-line assignment (RNA delivery)

In brief, whole-transcriptome libraries were processed with Parse split-pipe (v1.6.2) using Ensembl release 113 and analyzed with NVIDIA rapids-singlecell (v0.16.0)^22^. Genes detected in fewer than 100 cells were removed. Predicted doublets, cells with mitochondrial fraction greater than 20%, and additional outliers based on UMI counts, unique genes per cell, and top-gene fraction were excluded. Cell-line assignments were generated using scANVI^23^ with

Tahoe-100M-derived reference information and genotype-aware demultiplexing support from the Tahoe team. Low-confidence or missing assignments were refined using k-nearest-neighbor assignment to nearby high-confidence cells; cells unassigned after this step were excluded from downstream analysis.

### Whole transcriptome processing and cell-line assignment (plasmid delivery)

Whole-transcriptome and custom synthetic mRNA libraries were processed using 10x Genomics cellranger (v10.1.0)^24^ with the GRCh38-2024-A reference and analyzed with NVIDIA rapids-singlecell (v0.16.0). Whole transcriptome data were analyzed similarly to the RNA delivery method as described above, using rapids-singlecell for single-cell analysis and cell-line assignments generated with scANVI (v1.5.0) using Tahoe pool reference gene expression information from the RNA delivery dataset.

### Synthetic gene quantification and normalization

Synthetic RNA-derived reads were assigned to library elements and collapsed by cell line, sample, time point, and biological replicate. Concentration spike-ins were used to evaluate whether expected relative abundance ratios were recovered, and positive and negative stability controls were used to assess time-resolved recovery of known stable and destabilized sequence elements.

For episomal/plasmid experiments, fragment-level RNA counts were normalized to fragment-level DNA concentration in the input pool. This produced a DNA-normalized expression estimate for each fragment in each cell-line context. These values were analyzed alongside RNA-delivery measurements to separate transcriptional output from RNA stability and other post-transcriptional effects.

### Decay-rate estimation

For each library element and cell-line context, normalized abundance was calculated based on the number of copies per µL for each sample and modeled over time using (S_t = S_0 e^{-kt}), where (k) is the decay-rate parameter. Decay rates were estimated using all time points and in a sensitivity analysis excluding the 0-hour time point.

### Data Release

Both datasets focus on final, modeling-ready regulatory outputs rather than all intermediate processing artifacts. The release will include:

#### *Penta-47×27K* (5′ UTR / Episomal DNA)

- Final DNA-normalized expression estimates by library element and cell-line context.

#### *Tria-47×28K* (3′ UTR / Synthetic RNA)

- Final decay-rate matrices by library element and cell-line context.

## References

1. Zhang, J. et al. *Tahoe-100M*: A Giga-Scale Single-Cell Perturbation Atlas for Context-Dependent Gene Function and Cellular Modeling. Preprint at 10.1101/2025.02.20.639398 (2025).

2. Adamson, B. et al. A Multiplexed Single-Cell CRISPR Screening Platform Enables Systematic Dissection of the Unfolded Protein Response. Cell 167, 1867–1882.e21 (2016).

3. Dixit, A. et al. Perturb-Seq: Dissecting Molecular Circuits with Scalable Single-Cell RNA Profiling of Pooled Genetic Screens. Cell 167, 1853–1866.e17 (2016).

4. Replogle, J. M. et al. Mapping information-rich genotype-phenotype landscapes with genome-scale Perturb-seq. Cell 185, 2559–2575.e28 (2022).

5. Norman, T. M. et al. Exploring genetic interaction manifolds constructed from rich single-cell phenotypes. Science 365, 786–793 (2019).

6. Lotfollahi, M., Wolf, F. A. & Theis, F. J. scGen predicts single-cell perturbation responses. Nat. Methods 16, 715–721 (2019).

7. Roohani, Y., Huang, K. & Leskovec, J. Predicting transcriptional outcomes of novel multigene perturbations with GEARS. Nat. Biotechnol. 42, 927–935 (2024).

8. Theodoris, C. V. et al. Transfer learning enables predictions in network biology. Nature 618, 616–624 (2023).

9. Cui, H. et al. scGPT: toward building a foundation model for single-cell multi-omics using generative AI. Nat. Methods 21, 1470–1480 (2024).

10. Adduri, A. K. et al. Predicting cellular responses to perturbation across diverse contexts with State. Preprint at 10.1101/2025.06.26.661135 (2025).

11. Dong, M. et al. Stack: In-Context Learning of Single-Cell Biology. Preprint at 10.64898/2026.01.09.698608 (2026).

12. Wang, C. et al. X-Cell: Scaling Causal Perturbation Prediction Across Diverse Cellular Contexts via Diffusion Language Models. Preprint at 10.64898/2026.03.18.712807 (2026).

13. Svensson, V. et al. Back to basics: Observed statistics are sufficient to predict drug responses. 2026.06.09.731197 Preprint at 10.64898/2026.06.09.731197 (2026).

14. Hong, C. K. Y., Feng, F., Ramanathan, V., Liu, J. & Hansen, A. S. Genome structure mapping with high-resolution 3D genomics and deep learning. Preprint at 10.1101/2025.05.06.650874 (2025).

15. Wu, J., Wan, C., Ji, Z., Zhou, Y. & Hou, W. EpiFoundation: A Foundation Model for Single-Cell ATAC-seq via Peak-to-Gene Alignment. Preprint at 10.1101/2025.02.05.636688 (2025).

16. Siegel, D. A., Le Tonqueze, O., Biton, A., Zaitlen, N. & Erle, D. J. Massively parallel analysis of human 3′ UTRs reveals that AU-rich element length and registration predict mRNA destabilization. G3 GenesGenomesGenetics 12, jkab404 (2022).

17. Zhao, W. et al. Massively parallel functional annotation of 3′ untranslated regions. Nat. Biotechnol. 32, 387–391 (2014).

18. Franks, A., Airoldi, E. & Slavov, N. Post-transcriptional regulation across human tissues. PLOS Comput. Biol. 13, e1005535 (2017).

19. Gordon, M. G. et al. lentiMPRA and MPRAflow for high-throughput functional characterization of gene regulatory elements. Nat. Protoc. 15, 2387–2412 (2020).

20. Arnold, C. D. et al. Genome-wide assessment of sequence-intrinsic enhancer responsiveness at single-base-pair resolution. Nat. Biotechnol. 35, 136–144 (2017).

21. Lim, Y. et al. Multiplexed functional genomic analysis of 5’ untranslated region mutations across the spectrum of prostate cancer. Nat. Commun. 12, 4217 (2021).

22. Dicks, S., et al. GPU-accelerated single-cell analysis at scale with rapids-singlecell. Preprint at 10.48550/arXiv.2603.02402 (2026).

23. Xu, C. et al. Probabilistic harmonization and annotation of single-cell transcriptomics data with deep generative models. Mol. Syst. Biol. 17, MSB20209620 (2021).

24. Zheng, G. X. Y. et al. Massively parallel digital transcriptional profiling of single cells. Nat. Commun. 8, 14049 (2017).

25. Oikonomou, P., Goodarzi, H. & Tavazoie, S. Systematic Identification of Regulatory Elements in Conserved 3′ UTRs of Human Transcripts. Cell Rep. 7, 281–292 (2014).

26. Agarwal, V. et al. Massively parallel characterization of transcriptional regulatory elements. Nature 639, 411–420 (2025).

27. Khoroshkin, M. et al. A generative framework for enhanced cell-type specificity in rationally designed mRNAs. Preprint at 10.1101/2024.12.31.630783 (2024).

28. Reches, A., Berhani, O. & Mandelboim, O. A Unique Regulation Region in the 3′ UTR of HLA-G with a Promising Potential. Int. J. Mol. Sci. 21, 900 (2020).

29. Barreau, C. AU-rich elements and associated factors: are there unifying principles? Nucleic Acids Res. 33, 7138–7150 (2005).

30. Van Etten, J. et al. Human Pumilio Proteins Recruit Multiple Deadenylases to Efficiently Repress Messenger RNAs. J. Biol. Chem. 287, 36370–36383 (2012).

31. Wu, L., Fan, J. & Belasco, J. G. MicroRNAs direct rapid deadenylation of mRNA. Proc. Natl. Acad. Sci. 103, 4034–4039 (2006).

32. Lupini, L. et al. miR-221 affects multiple cancer pathways by modulating the level of hundreds messenger RNAs. Front. Genet. 4, (2013).

33. Grimson, A. et al. MicroRNA Targeting Specificity in Mammals: Determinants beyond Seed Pairing. Mol. Cell 27, 91–105 (2007).

34. Rosado-Tristani, D. A. et al. CisBP-RNA: a web resource for eukaryotic RNA-binding proteins and their motifs. Nucleic Acids Res. 54, D98–D105 (2026).

35. Agarwal, V. & Shendure, J. Predicting mRNA Abundance Directly from Genomic Sequence Using Deep Convolutional Neural Networks. Cell Rep. 31, 107663 (2020).

36. Hao, M. et al. Large-scale foundation model on single-cell transcriptomics. Nat. Methods 21, 1481–1491 (2024).

37. Penzar, D. et al. LegNet: a best-in-class deep learning model for short DNA regulatory regions. Bioinformatics 39, btad457 (2023).

38. Hunter, T. L. et al. In vivo CAR T cell generation to treat cancer and autoimmune disease. Science 388, 1311–1317 (2025).

39. Leppek, K. et al. Combinatorial optimization of mRNA structure, stability, and translation for RNA-based therapeutics. Nat. Commun. 13, 1536 (2022).

40. Jung, S.-J. et al. RNA stability enhancers for durable base-modified mRNA therapeutics. Nat. Biotechnol. https://doi.org/10.1038/s41587-025-02891-7 (2025) doi:10.1038/s41587-025-02891-7.

